# Morphodynamic domains enable integration of live morphometrics and spatial transcriptomics

**DOI:** 10.64898/2026.08.13.744578

**Authors:** Adrien Leroy, Eric van Leen, Maria Balakireva, Lale Alpar, Fabian Gärtner, Isabelle Gaugué, Stéphane Pelletier, Rémi Pigache, Mehdi Ech-Chouini, Julien Delpierre, Stéphane Rigaud, Floris Bosveld, Lorette Noiret, Yohanns Bellaïche

**Affiliations:** Institut Curie, PSL Research University, CNRS UMR3215, INSERM U934, F-75248 Paris Cedex 05, France; Sorbonne Université, CNRS UMR3215, INSERM U934, F-75005, France

## Abstract

Tissue development emerges from the coordinated behaviors of thousands of cells, orchestrated by gene regulatory networks. Recent methodological advances now enable high-resolution live imaging of cell- and tissue-scale dynamics and the construction of spatially resolved gene expression atlases. However, quantitatively linking these modalities remains a central challenge, limiting our ability to understand how gene regulatory networks drive cell- and tissue-scale behaviors. Here, using the *Drosophila* thorax epithelium as a model system, we introduce an analytical and computational framework based on tissue morphodynamic domains: regions defined by coherent cell and tissue dynamics extracted from live imaging morphometrics. Integrating morphodynamic domains with spatial transcriptomics enables the inference of gene regulatory networks associated with distinct spatial cell- and tissue-level behaviors. Statistical cross-scale analyses further allow the interrogation and validation of gene function, confirming or revealing regulators of specific morphogenetic dynamics. In particular, our framework uncovers a role for the Toll-like receptor Tollo in modulating tissue flow, contraction, and apoptosis. Together, our work establishes a framework that integrates spatial transcriptomics with quantitative, multiscale live morphometrics, providing a generalizable strategy to probe and understand developmental processes.

## Introduction

Development entails the formation of reproducible tissue organizations and shapes controlled by gene expression.^1^ While conserved regulators of tissue development have been uncovered, recent advances in live imaging and spatial transcriptomics are poised to drastically improve our understanding of tissue development by identifying gene regulatory networks (GRNs) governing cell and tissue dynamics. Yet the multimodal integration of quantitative live imaging and spatial transcriptomic datasets remains a major challenge.

Advanced live imaging has been instrumental in illuminating the complexity of collective cell and tissue dynamics during development.^2^ Complementarily, multiscale formalisms - quantifying cell shape changes, divisions, apoptosis and cell-cell rearrangements - have enabled comprehensive morphometric characterization of cell- and tissue-level dynamics in space and time.^3–9^ Such high-dimensional quantitative datasets have provided valuable insights into the mechanisms driving collective cell behaviors and large-scale tissue movements.^10–14^ Complementing these advances, single-cell RNA sequencing (scRNA-seq) and spatial transcriptomics have recently transformed our ability to profile gene expression within tissues.^15–18^ In particular, computational algorithms that reconstruct gene expression maps from scRNA-seq data now enable the description of spatial transcriptomes and the inference of GRNs, even in the absence of direct spatial measurements.^19–38^ More recent approaches have further enabled the reconstruction of high-resolution transcriptome profiles of tissues, organs, and entire organisms across successive developmental stages.^39–45^ By correlating gene expression patterns between developmental stages, computational methods have been used to infer coarse-grained cell reorganization at successive time points, deducing how gene expression patterns evolve with global cell displacements.^43,45–48^ However, correlating gene expression across time does not directly capture cell dynamics.^49^ As a result, these approaches require validation with live-imaging and remain to be complemented by comprehensive live morphometrics of cell- and tissue-scale dynamics to infer the function of candidate regulators. Moreover, establishing causal links between candidate regulators and cell and tissue dynamics requires experimental perturbations that directly test gene function.

Integrating quantitative live morphometrics with spatially resolved gene expression and validating this integration through loss-of-function perturbations pose several challenges. First, cell and tissue dynamics are captured using a variety of quantitative descriptors including scalars, vectors, and tensors.^3–5,7,8,50–56^ These quantities must then be related to local gene expression, which is typically measured as a scalar value. Second, tissue development is a continuous, high-dimensional process, calling for strategies to reduce its dimensionality to facilitate the quantitative description and exploration of cell and tissue behaviors across space and time.^56–58^ Third, mechanical interactions within tissues are inherently non-local.^14,59^ Consequently, a local gene perturbation that modulates tissue mechanics can produce effects at distant sites and during later developmental stages. Therefore, there is a need for objective analytical methods to identify both local and non-local effects of gene perturbations. To date, no framework exists to navigate this complexity while simultaneously addressing these challenges to investigate the mechanisms driving cell and tissue dynamics during development.

Here, we developed a computational framework to integrate live imaging data with spatial transcriptomics, enabling the discovery of GRNs governing tissue development. Our approach leverages millions of tissue-scale live cell imaging morphometrics together with spatial transcriptomics in the *Drosophila* pupal dorsal thorax (notum) epithelium. Central to this framework is the definition of morphodynamic domains, regions characterized by coherent combinations of cell- and tissue-level behaviors. By mapping spatial transcriptomic information onto these domains, we identify candidate GRNs associated with distinct morphogenetic dynamics. The framework recapitulates known regulators of epithelial morphogenesis and uncovers a previously uncharacterized role for the Toll-like receptor Tollo in modulating tissue flow, contraction, and apoptosis. Together, our work provides both a generalizable analytical strategy and a resource for bridging high-dimensional live morphometrics with spatial transcriptomics to investigate the gene regulatory control of tissue development.

## Results

### Multiscale clustering of cell and tissue dynamics to define morphodynamic domains

The *Drosophila* pupal dorsal thorax (notum) epithelium is a powerful model system for studying conserved mechanisms controlling tissue dynamics and cell fate specification (Figures 1A and 1B).^4,6,60–72^ As observed in multiple tissues, its morphogenesis relies on complex spatiotemporal coordination of morphodynamic behaviors across its ∼10^4^ cells, including tissue deformation and flow, driven by distinct cellular dynamics (Video S1).^4,6,73–76^ This includes regions undergoing oriented cell divisions, rearrangements, apoptosis, and shape changes that collectively shape the tissue.^4,6,73–76^ Its inherent spatiotemporal heterogeneity makes it a suitable model to unravel the gene regulatory landscapes governing diverse morphogenetic programs via spatial transcriptomics.

**Figure 1.**
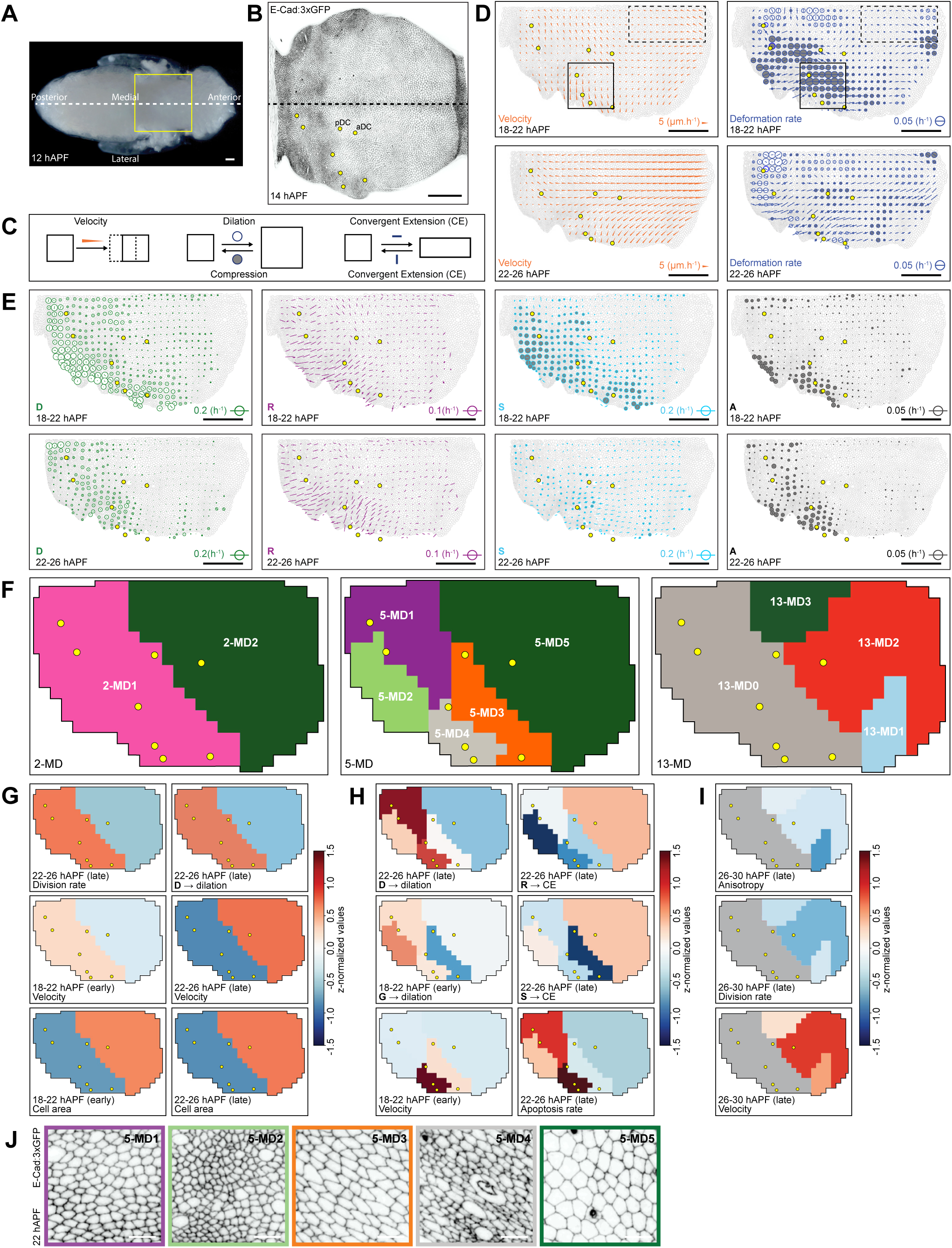
Characterization of morphodynamic domains during notum morphogenesis. (**A**) Image of a *Drosophila* pupa without pupal case at 12 hAPF. Yellow box: region shown in B; white dashed line: midline (axis of left-right symmetry). (**B**) Confocal image of E-Cad:3xGFP in the notum epithelium at 14 hAPF. Black dashed line: midline; yellow dots: landmark macrochaetae in the right hemi-notum. Posterior and anterior dorsocentral macrochaetae (pDC and aDC) are indicated. (**C**) Schematic illustrating how tissue flow (left), isotropic tissue dilatation and contraction (middle), as well as convergent extension (CE) (right) contribute to tissue deformation. Orange arrowhead: tissue flow velocity; white circle: magnitude of isotropic dilatation; grey circle: magnitude of isotropic contraction; blue bars: orientation and magnitude of anisotropic convergent extension. (**D**) Maps of the average (*N* = 8) tissue flow velocity (left) and tissue deformation rate (right) in the right hemi-notum epithelium, averaged over 18-22 hAPF (top) and 22-26 hAPF (bottom), and plotted onto a reference animal. The boxes (at 18-22 hAPF) indicate the region of high lateral tissue flow and contraction (solid line) as well as the region exhibiting high posterior-to-anterior tissue flow in absence of deformation (dashed line). Orange arrows indicate the direction and amplitude of tissue flow. Grey circles indicate the amplitude of isotropic contraction, and white circles indicate the amplitude of isotropic dilation, while bars indicate the orientation and anisotropy of CE rate. Yellow dots: landmark macrochaetae. (**E**) Maps of the average (*N* = 8) contribution of cell division **D**, cell rearrangement **R**, cell shape change **S**, and apoptosis **A** to notum tissue deformation, averaged over 18-22 hAPF (top) and 22-26 hAPF (bottom), and plotted onto a reference animal. Grey circles indicate the amplitude of isotropic contraction, and white circles indicate the amplitude of isotropic dilation, while bars indicate the orientation and anisotropy of CE rate. Yellow dots: landmark macrochaetae. (**F**) Spatial maps of morphodynamic domains (MD) for 2 clusters (2-MD, left), 5 clusters (5-MD, middle), and 13 clusters (13-MD, right) inferred from the average of *N* = 8 hemi-nota live movies. For the 13-MD, only the anterior domain resolved into three sub-clusters is shown, whereas the posterior domain is shown in grey as a single aggregated cluster. Note, 13-MD0 is identical to 2-MD1. Similarly, 2-MD2 is identical to 5-MD5. Yellow dots: landmark macrochaetae. (**G**) Maps of the z-normalized values for 2-MD for the tissue normalized division rate in the late phase (top left), the division contribution to tissue isotropic dilatation in the late phase (**D** → dilation, top right), tissue velocity in the early (middle left) and in the late phase (middle right) as well as cell area in the early (bottom left) and in the late phase (bottom right). Yellow dots: landmark macrochaetae. (**H**) Maps of the z-normalized values for 5-MD for the contribution of divisions to tissue dilatation rate in the late phase (**D** → dilation, top left), the contribution of cell rearrangement to tissue convergent extension rate in the late phase (**R** → CE, top right), the tissue isotropic dilation rate in the early phase (**G** → dilation, middle left), the cell shape change contribution to tissue convergent extension rate in the late phase (**S** → CE, middle right), the tissue velocity in the early phase (bottom left), and the tissue normalized apoptosis rate in the late phase (bottom right). (**I**) Maps of the z-normalized values for 13-MD for cell shape anisotropy in the late phase (top), the tissue normalized division rate in the late phase (middle), and the tissue velocity in the late phase (bottom). The grey domain (13-MD0, identical to 2-MD1) represents an aggregation of domains that were excluded for the interpretation. Yellow dots: landmark macrochaetae. (**J**) Confocal images of E-Cad:3xGFP in 5-MD1 to 5-MD5 at 22 hAPF highlighting differences in cell area and anisotropy in each morphodynamic domain. Scale bars: 100 µm (A, B, D, E), 10 µm (J); velocity and deformation rate are indicated (D, E).

To build an analytical and computational framework for integrating live imaging with spatial transcriptomics, we hypothesized that a critical step would be to organize tissues into domains characterized by coherent dynamic cell behaviors, hereafter referred to as ‘morphodynamic domains’. Such tissue segmentation enables both a quantitative description of tissue morphogenesis and the identification of domains within which domain-specific GRNs, associated with similar collective cell dynamics, can be inferred. To delineate the tissue morphodynamic domains, we first analyzed global tissue flows and deformations to quantitatively define distinct temporal phases of morphogenesis. We recorded 8 control time-lapse movies of notum tissues expressing the apical adherens junction (AJ) marker E-Cad:3xGFP at 5 min intervals, capturing over 12 h of tissue development. As previously described,^4,6^ we quantitatively measured tissue flow and deformation using particle image velocimetry (PIV) and subsequently averaged these measurements across the 8 movies after spatiotemporal alignment. Because the notum is bilaterally symmetric, the measurements are thereafter plotted on the right hemi-notum. Flow and deformation were measured in 40 µm x 40 µm spatial bins, ensuring robust measurements, while retaining sufficient resolution to assess how local cell dynamics contribute to global tissue morphogenesis.^4,6^ Extending our previous findings, analysis of the average velocity magnitude profiles across the tissue revealed three distinct developmental phases between 14 and 26 hours after puparium formation (hAPF) (Figure S1A; Video S1). The initial phase, spanning 14-18 hAPF, is characterized by low overall tissue flow and minimal tissue deformation (Figure S1A).^4,6^ The second phase, from 18-22 hAPF, shows a marked increase in tissue flow, including the emergence of a major lateromedial flow pattern accompanied by extensive lateral contraction (Figures 1C, 1D and S1A). Finally, the third phase, 22-26 hAPF, exhibits strong posterior-to-anterior tissue flow, most pronounced in the anterior medial region, where tissue deformation is minimal (Figures 1D and S1A). Thus, as previously performed in multiple tissues, the continuous process of notum development can be subdivided into successive phases.

Given the significant morphogenetic events during the latter two phases (18-22 hAPF and 22-26 hAPF), we focused our subsequent analyses on characterizing the spatiotemporal cell dynamics underlying tissue flows and deformations in these intervals. Cells from the 8 time-lapse movies were segmented and tracked, followed by quantitative analysis of cell and tissue dynamics, yielding millions of measurements (see Methods). This included measurements using a previously validated formalism that decomposes local tissue deformation rates into distinct cellular processes, cell division, cell rearrangement, apoptosis, and cell shape changes, to quantify the contribution of each process to overall tissue deformation.^4,6^ We also measured static features (cell area and anisotropy), dynamic metrics (division and apoptosis rates), as well as velocity flows and deformation rates. Altogether, this yielded a dataset of 15 features averaged over the two developmental phases, totaling 30 measured quantities across the tissue, capturing the diversity and heterogeneity of cell and tissue dynamics (Figures 1D and 1E; Video S2; Table S1).

To investigate how these 30 measured quantities relate to the spatiotemporal cell and tissue dynamics during notum morphogenesis, we first applied principal component analysis (PCA) for dimensionality reduction, mitigating potential bias due to redundancy among the measured features. We then retained the first 14 principal components (PCs), which collectively accounted for 90% of the total variance, and used them to cluster tissue dynamics into morphodynamic domains (Figures S1B-S1D). To automatically define morphodynamic domains, we used spatially constrained agglomerative clustering.^77^ This approach extends standard hierarchical clustering by incorporating a spatial connectivity constraint,^78^ ensuring that only spatially adjacent data points are merged.^79^ This method is particularly well suited for spatial data, as it respects the geometric relationships between points and allows for the identification of contiguous regions. Importantly, it provides a key advantage over previously implemented methods^58^ by enabling exploration of morphogenesis through a ‘zooming in’ approach, whereby the tissue can be segmented sequentially into smaller nested domains depending on the chosen resolution for analyzing its dynamics. To quantitatively guide our selection of the number of domains, we employed silhouette analysis,^80^ which evaluates how well morphodynamic domains are separated. We calculated the average silhouette score for domain numbers ranging from 2 to 13 (Figure S1E). The analysis revealed a maximum at 5 domains (score = 0.295), indicating this as the statistically optimal level of granularity (Figures S1E and S1F). We therefore focused our analysis on three levels of clustering: a coarse 2-domains partition (2-MD), corresponding to the primary anteroposterior dichotomy of the tissue; the statistically optimal 5-domains partition (5-MD), as indicated by the silhouette score; and a finer 13-domains partition (13- MD), which further resolved the anterior domain into three subdomains for more detailed analysis (Figure 1F).

This multi-level segmentation of tissue morphogenesis enables the description of tissue dynamics at different resolutions by identifying combinations of cell behaviors that define each morphodynamic domain. These can be represented as heat maps to highlight the combination of features characterizing each domain (Figures S1G-S1I), or as tissue maps showing the spatial distribution of features associated with tissue morphogenesis (Figures 1G-1I). At the broadest level, a 2-domain separation divides the notum into a posterior (2- MD1) and an anterior (2-MD2) domain based on several quantitative features (Figures 1F and S1G). In particular, the posterior domain exhibits a higher division rate and a stronger contribution of cell division to tissue dilation during the late phase (Figures 1G and S1G). The anterior domain is characterized by lower tissue velocity in the early phase and higher velocity in the late phase, as well as larger cell area throughout both phases (Figures 1G and S1G). Increasing the granularity to 5 domains subdivides the 2-MD1 posterior domain into 4 subdomains (Figure 1F). Among these subdomains, 5-MD1 is characterized by increased dilation associated with cell division during the late phase, whereas 5-MD2 exhibits the lowest contribution of cell shape changes to tissue dilation in the early phase, along with lower velocity and a reduced contribution of cell rearrangements to tissue convergent extension in the late phase (Figures 1H and S1H). The remaining posterior clusters are distinguished by distinct quantitative features: 5-MD3 shows lower dilation during the early phase and a reduced contribution of cell shape changes to convergent extension in the late phase (Figures 1H and S1H), whereas 5-MD4 exhibits the highest tissue velocity in the early phase, the highest apoptosis rate in the late phase (Figure 1H), and the most anisotropic cells across both phases (Figures 1J and S1H). This partitioning illustrates that the posterior region of the tissue contains multiple, distinct subdomains characterized by specific cell and tissue dynamics. Finally, further subdivision into 13 domains separates the anterior 2- MD2 domain (also corresponding to the 5-MD5) into three subdomains (Figure 1F). 13-MD1 displays low cell shape anisotropy during both phases; 13-MD2 is characterized by higher flow velocity and a lower division rate in the late phase; and 13-MD3 shows strong tissue convergent extension during the late phase (Figures 1I and S1I). Together, this clustering approach enables delineating distinct morphodynamic domains based on a large set of static and dynamic measurements and to extract their main morphodynamic features.

### Reconstruction of the spatial transcriptome of the notum prior to tissue morphogenesis

To identify GRNs associated with distinct morphodynamic domains, we first implemented a method to reconstruct the spatial transcriptome of the developing notum by mapping scRNA-seq data onto a reference gene pattern atlas (Figure 2A). We performed scRNA-seq on 31,991 cells derived from over 160 dissociated nota at 15 hAPF, a timepoint preceding any significant morphogenetic events (Figure S1A).^4,6^ After analyzing the scRNA-seq data using Seurat^81^ and performing quality control and filtering, we retained 12,187 high-quality cells and 9,699 genes, with a median of 26,142 unique molecular identifiers (UMIs) per cell. Within our dataset, we identified several populations corresponding to cell types known to be present within the notum tissue or associated with it: notum epithelial cells (*n* = 7,610 cells); neck epithelial cells (*n* = 434 cells); neuronal sensory organ precursors (*n* = 174 cells); myoblasts and tracheal cells (*n* = 2,561 cells); and an unannotated cell cluster of unknown origin (*n* = 1,408 cells) (Figures 2A and S2A). Complementarily, we constructed an atlas of 24 gene patterns, primarily using available fluorescently tagged reporters and generating fluorescent CRISPR knock-in reporters for 11 genes (see Methods). For each gene, patterns from at least 3 animals were averaged to generate the atlas. Using well-defined macrochaetae landmarks (Figure 1B), our pattern rescaling and registration algorithm can average these patterns to 15 µm × 15 µm bins, corresponding to a resolution of 1.8 ± 0.7 cells (mean ± SD). Here, we discretized the 24 atlas patterns into 40 µm × 40 µm bins (12.5 ± 5.2 cells, mean ± SD) to explore the link between gene expression and morphogenesis, because this approach: (i) aligns with the spatial footprint of the morphometric measurements; (ii) increases counts per grid position, thereby improving the signal-to-noise ratio; and (iii) enables subsequent statistical comparisons between transcriptomic and morphodynamic datasets. To map each sequenced single epithelial cell, we developed a simple mapping method named ‘virtual Gene Expression Pattern’ (vGEP). For each spatial bin in the grid, we calculated the correlation coefficient between the local expression profile of the 24 atlas patterns and the expression profile of each single cell. We then reconstructed the transcriptome at each grid position by computing a weighted average of gene expression from the top-correlating cells, with weights proportional to correlation strength. Focusing on genes expressed in at least 50 cells to ensure statistical robustness, we reconstructed a spatial transcriptome encompassing the patterns of 4,312 genes within the notum epithelium (available at https://ybellaichelab.shinyapps.io/NotumvGEP/).

**Figure 2.**
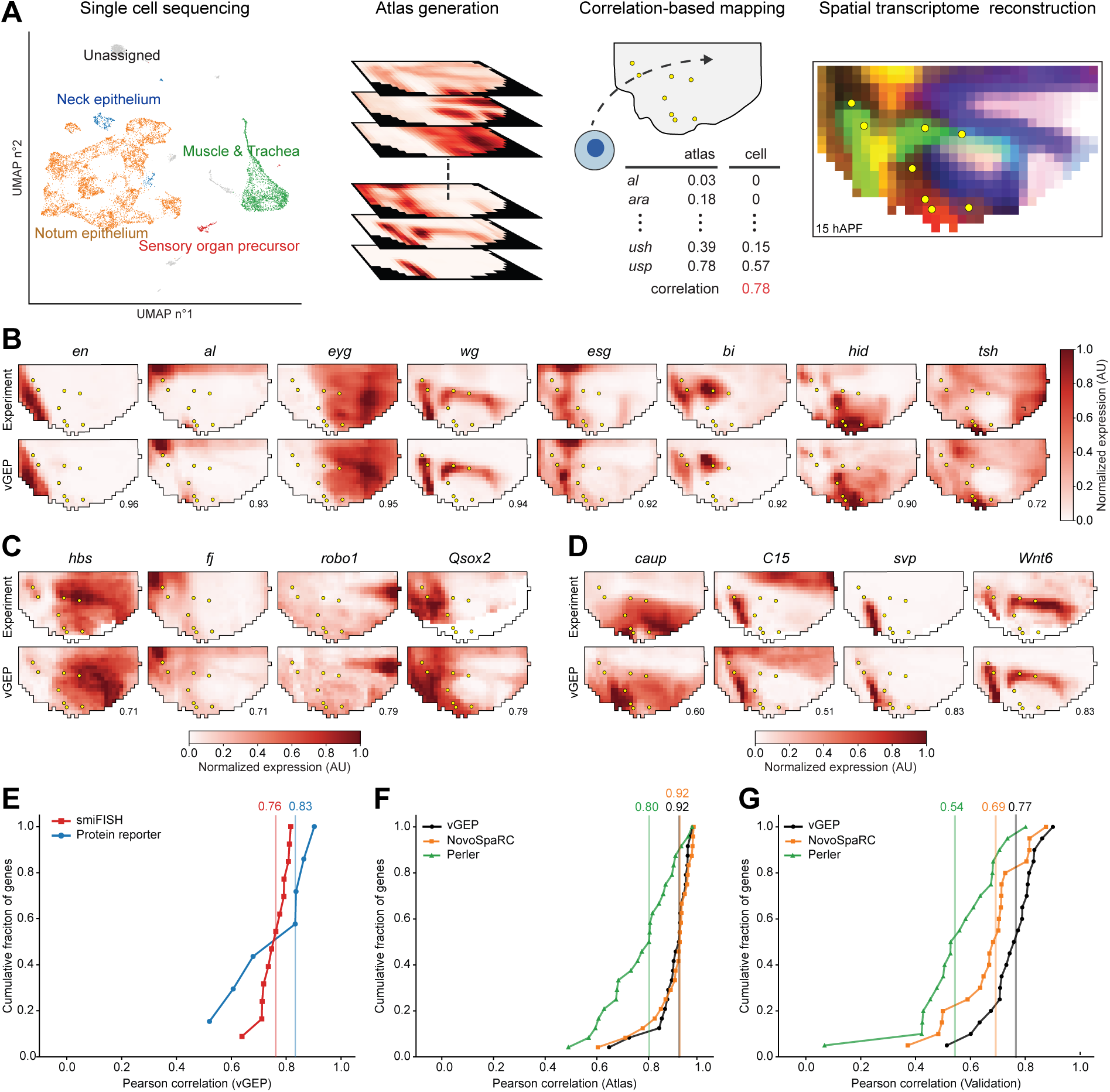
Reconstruction and validation of the notum spatial transcriptome. (**A**) Schematic of spatial transcriptome reconstruction using the virtual Gene Expression Pattern (vGEP) method. Single-cell RNA-seq data are first clustered by cell type (UMAP, left). Transcriptomic profiles of epithelial cells (cell vector) are correlated with local patterns of the 24 atlas genes (atlas vector, middle) to reconstruct the transcriptome for each spatial bin in the notum epithelium at 15 hAPF (right). Yellow dots: landmark macrochaetae. (**B**) Average experimental expression patterns and vGEP reconstructions of 8 representative atlas genes: *en*, *al*, *eyg*, *wg*, *esg*, *bi*, *hid*, and *tsh*. r values (Pearson correlation between experiment and vGEP) are indicated. Yellow dots: landmark macrochaetae. (**C**) Average experimental expression patterns and vGEP reconstructions of 4 representative smiFISH validation genes: *hbs*, *fj*, *robo1*, and *Qsox2*. r values (Pearson correlation between experiment and vGEP) are indicated. Yellow dots: landmark macrochaetae. (**D**) Average experimental expression patterns and vGEP reconstructions of 4 representative protein reporter genes: *caup*, *C15*, *svp*, and *Wnt6*. r values (Pearson correlation between experiment and vGEP) are indicated. Yellow dots: landmark macrochaetae. (**E**) Cumulative distribution of Pearson correlations between vGEP and 13 smiFISH validation expression patterns (r_median_ = 0.76) as well as 7 protein reporters and (r_median_ = 0.83). Each point on the curve represents the fraction of genes with a correlation at or below the corresponding value. (**F**) Cumulative distribution of Pearson correlations between predicted and reference (atlas) spatial expression profiles for the 24 atlas genes (training set) using vGEP (r_median_ = 0.92), NovoSpaRC (r_median_ = 0.92), and Perler (r_median_ = 0.77). Each point on the curve represents the fraction of genes with a correlation at or below the corresponding value. (**G**) Cumulative distribution of Pearson correlations between predicted and spatial expression profiles for 20 validation genes (combined from E) using vGEP (r_median_ = 0.77), NovoSpaRC (r_median_ = 0.69), and Perler (r_median_ = 0.54). Each point on the curve represents the fraction of genes with a correlation at or below the corresponding value.

To validate our spatially reconstructed transcriptome, we first selected 13 genes with diverse predicted spatial patterns that were not included in the gene pattern atlas and determined their expression using single-molecule fluorescence *in situ* hybridization (smiFISH).^82^ Comparing the predicted and experimental expression patterns for the 13 validation genes revealed the accuracy of our mapping method, with high correlation between vGEP and experimental spatial patterns (r_median_ = 0.76; Figures 2B-2E). This also provided *a posteriori* validation of using fluorescent reporter CRISPR knock-ins to construct our atlas patterns. We then analyzed 7 available fluorescent reporters and found similarly high correlations between vGEP and experimental spatial patterns (r_median_ = 0.83; Figures 2D and 2E). Finally, we benchmarked our method against two established spatial reconstruction algorithms: NovoSpaRC/Moscot^83^ and Perler^21^ (Figures 2F, 2G and S2B-S2F). All three methods performed well in reconstructing the atlas expression patterns (r_median_ = 0.92 for NovoSpaRC, 0.80 for Perler, and 0.92 for vGEP, Figures 2F, S2C and S2E). Importantly, vGEP showed improved predictive ability for gene expression patterns not used in the reconstruction. Specifically, vGEP achieved a median correlation of 0.77 with 20 validation genes, compared to 0.69 for NovoSpaRC and 0.54 for Perler (Figures 2G, S2D and S2F). This improved performance was achieved using only a single tunable parameter, the number of top-correlating cells used per bin, while drastically reducing computational requirements (Figure S2B; see Methods). Thus, vGEP provides an efficient, scalable approach for reconstructing spatial transcriptomes from large cell numbers.

### Analysis of GRNs associated with each morphodynamic domain

Focusing on the optimal 5-MD tissue segmentation and utilizing the spatial transcriptome, we subsequently explored the link between spatial gene expression and tissue dynamics by inferring GRNs operating within each of the morphodynamic domains. By thresholding the tau specificity score distribution of gene expression,^84,85^ we first identified a set of 38 transcription factors (TFs) that are maximally expressed within one of the 5 morphogenetic domains (Figures 3A and S3A). Second, to reconstruct domain-specific GRNs, we applied the GENIE3 framework^86^ to predict TF-target relationships based on non-linear co-expression in single cells of the notum epithelium, enabling the identification of both positively and negatively regulated targets. This analysis yielded 598 up- or downregulated target genes distributed across different morphodynamic domains (Figures 3B, 3C and S3B-3BD).

**Figure 3.**
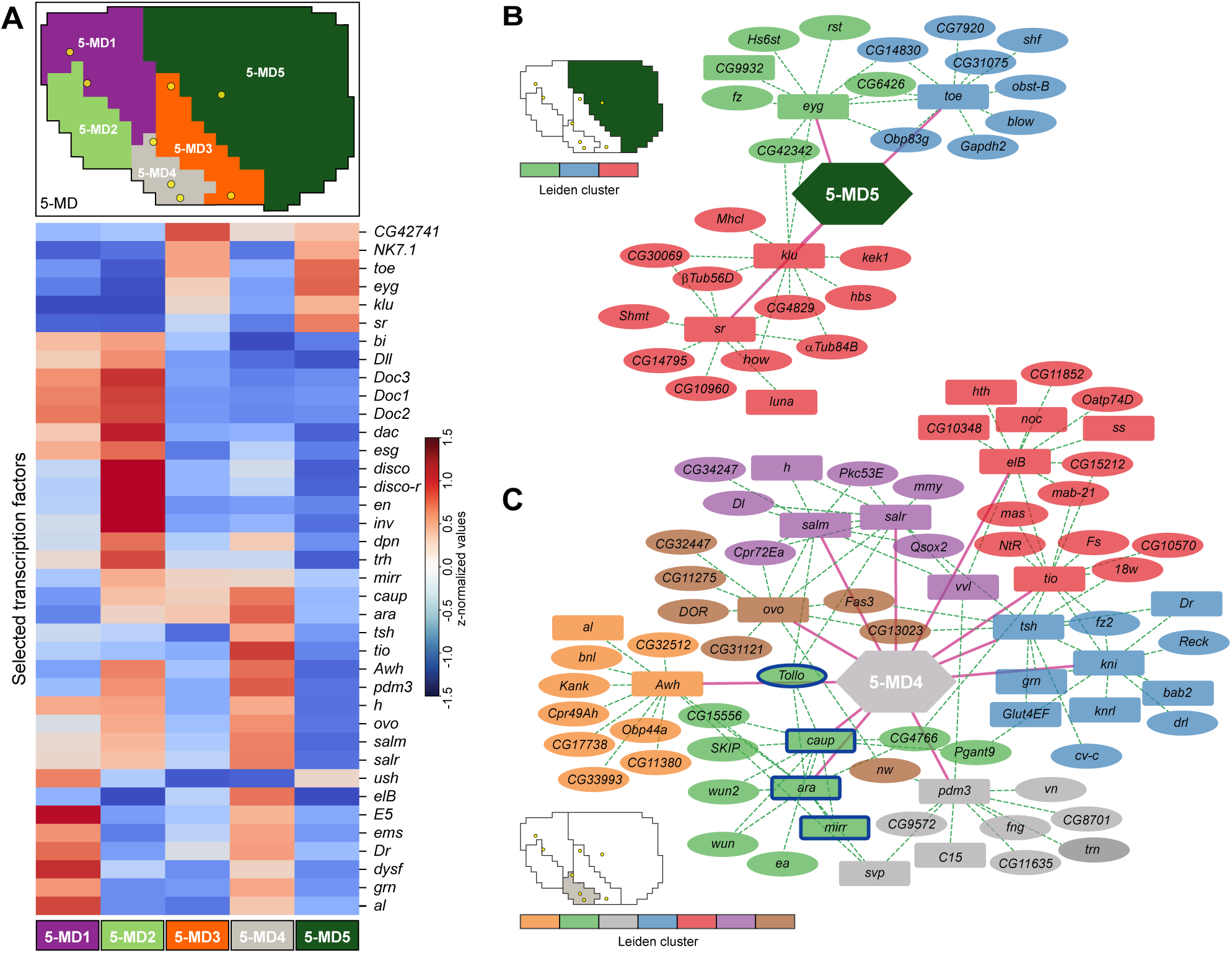
Linking morphodynamic domains to their underlying GRNs. (**A**) Z-normalized values of 38 selected transcription factors (TFs) identified as having domain-specific expression (top 10% tau score) across the 5-MD domains. The spatial map of 5-MD is indicated at the top. Yellow dots: landmark macrochaetae. (**B**) Inferred GRN for the anterior morphodynamic domain 5-MD5. Nodes represent TFs (rectangles) and their top 10 predicted target genes (ellipses). Colors indicate Leiden clusters. Pink solid lines indicate high specificity (τ > 0.36). Green dashed lines indicate putative regulation identified through GRN inference. (**C**) Inferred GRN for the lateral morphodynamic domain 5-MD4. Nodes represent TFs (rectangles) and their top 10 predicted target genes (ellipses). Colors indicate Leiden clusters. Pink solid lines indicate high specificity (τ > 0.36). Green dashed lines indicate putative regulation identified through GRN inference. Blue outlines highlight *Tollo* and its predicted upstream TFs belonging to the Iroquois complex genes (*caup*, *ara*, *mirr*).

Analysis of the GRNs revealed distinct transcriptional signatures associated with each morphodynamic domain. In the most anterior domain (5-MD5), characterized by high cell velocity and large apical areas (Figures 1F, 1G, 1J and S1H), we found specific expression of the TFs Stripe (Sr), Klu, Eyg, and Toe (Figure 3B). Network inference identified a regulatory module centered on Sr, consistent with our recent findings showing that Sr regulates anterior tissue flow through collective cell migration (Guirao et al., *submitted*). Interestingly, the finer 13-MD clustering revealed that *Sr* expression is specifically enriched in the 13-MD2 subdomain, which exhibits significantly higher tissue flow than adjacent regions (Figures 1G, S1I and S3E), further supporting its key role in regulating collective cell migration (Guirao et al., *submitted*). The most posterior and medial morphodynamic domain (5-MD1), which exhibits elevated dilation associated with cell division during the late phase (Figures 1H and S1H), is characterized by a distinct transcriptional landscape defined by the expression of Al, Ush, Dr, Dysf, Grn, and E5 (Figure S3B). Notably, our GRN analysis identified Four-jointed (Fj) as a specific target of Ush. Fj has previously been shown to regulate tissue flow in this region together with the protocadherin tumor-suppressors Fat and Dachsous, by controlling cell-cell junction tension and rearrangements.^6^ Morphodynamic domain 5-MD2 was associated with a broad set of TFs, including, Dac, H, Disco, Dorsocross genes (Doc1, Doc2, Doc3), Mamo, Dpn, Bi, Esg, Dll, and Trh (Figure S3C). In contrast, 5-MD3 contained fewer specific factors, primarily NK7.1 and the uncharacterized gene CG42741 (Figure S3D). Finally, we focused on the lateral domain (5-MD4), which displays prominent tissue flow velocity during the early phase as well as high apoptosis rate in the late phase (Figures 1F, 1H and S1H). The TF regulatory landscape within this morphodynamic domain included Awh, Pdm3, Ovo, Salm, ElB, Tio as well as the Iroquois complex genes (Ara, Caup, Mirr) (Figure 3B). Interestingly, the inferred GRN for this domain included Toll-like receptor (TLR) signaling components as targets, including Ea and, notably, the TLR Tollo (Figures 3B and S3F). This enrichment is particularly intriguing, as previous studies have implicated TLRs in promoting tissue flow through cell rearrangements and tissue invagination.^87–91^ Finally, gene expression analysis of the putative Tollo ligand, Spz3^92^ revealed that it is also predicted to be highly expressed in 5-MD4 (Figure S3F).

Together, these analyses show that integrating live imaging with spatial transcriptomics via morphodynamic domains enables the identification of candidate regulators of tissue dynamics. To functionally validate our framework, we next investigated the role of Tollo in epithelial tissue morphogenesis.

### Characterizing the contribution of Tollo to notum epithelial morphogenesis

To investigate the role of Tollo, we first characterized its localization using a Tollo:GFP knock-in transgene. This confirmed that Tollo was enriched in 5-MD4, forming a lateral to medial gradient that extends into 5-MD3 (Figures 4A and S3F). Our GRN analyses predicted that Tollo expression is positively controlled by the three TFs of the Iroquois complex, Ara, Caup and Mirr (Figure 3C). These three TFs often function redundantly.^93–97^ While disrupting Ara and Caup function using RNAi did not affect Tollo:GFP localization (not shown), simultaneous inhibition of Ara, Caup and Mirr (*ara,caup,mirr^RNAi^*) abrogated the Tollo:GFP signal, confirming the prediction of our GRN analysis (Figure 4B). We then tested whether Tollo controls notum morphogenesis by knocking-down Tollo activity using a dsRNA against Tollo (*Tollo^RNAi^*) and quantifying tissue flow and deformation from time-lapse E-Cad:3xGFP movies. Tissue flows and deformations in *Tollo^RNAi^* (*N* = 5) were averaged and subsequently subtracted from those of *v^RNAi^* control animals (*N* = 8) (*Tollo^RNAi^* - *v^RNAi^*) to determine spatiotemporal differences in velocity and deformation. This approach revealed differences in both velocity and deformation profiles between *Tollo^RNAi^* and control tissues (Video S3). These differences were heterogeneously distributed across the tissue and varied over time, posing a challenge for objectively defining where and when Tollo loss-of-function impacts epithelial morphogenesis. We therefore sought to develop a general approach to identify significant spatiotemporal morphodynamic changes between experimental conditions.

**Figure 4.**
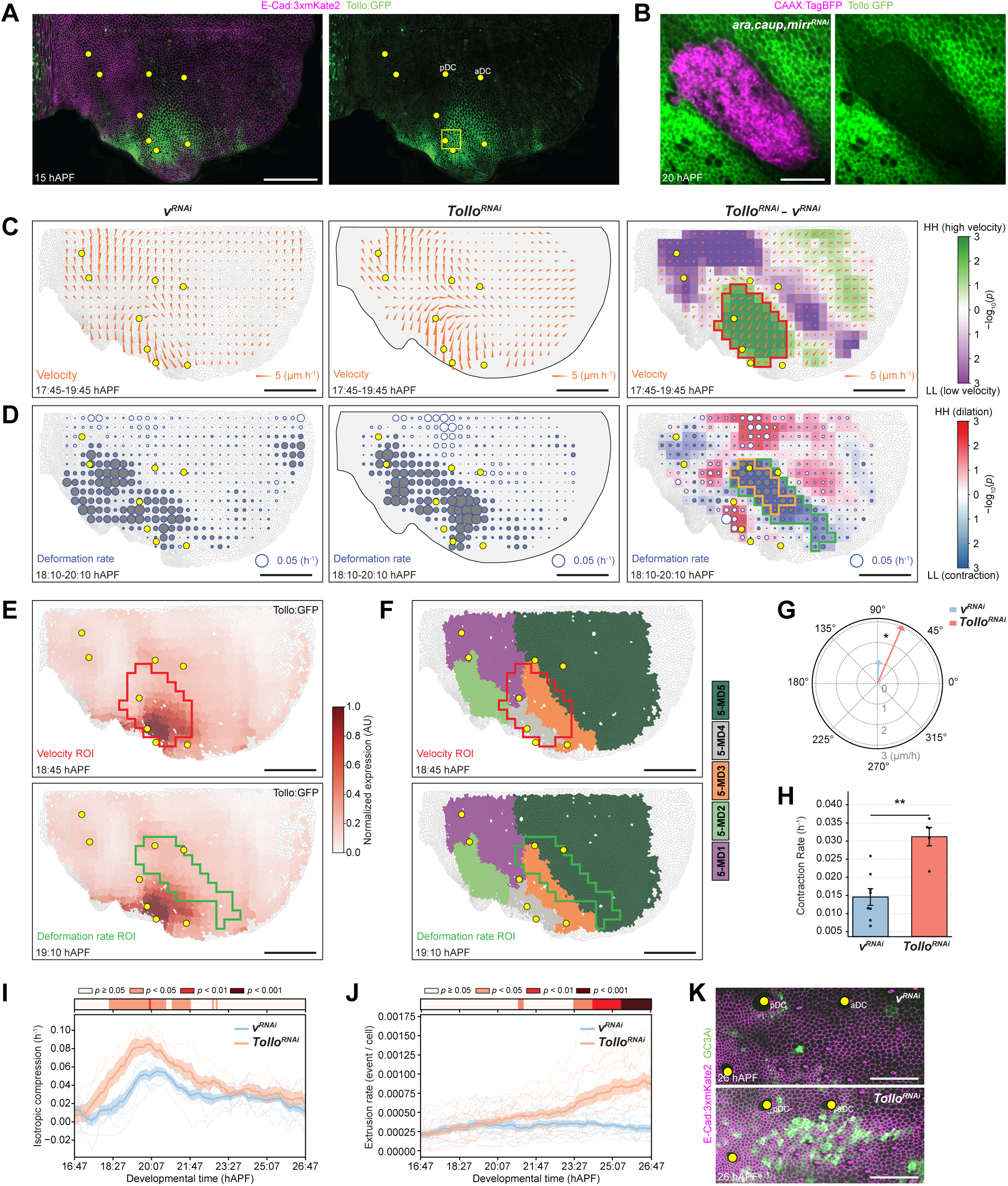
Spatiotemporal morphodynamic changes in *Tollo^RNAi^* tissues. (**A**) Confocal images of E-Cad:3xmKate2 and Tollo:GFP in the notum epithelium at 15 hAPF. Yellow box: region analyzed in B. Yellow dots: landmark macrochaetae. Posterior and anterior dorsocentral macrochaetae (pDC and aDC) are indicated. (**B**) Confocal image of Tollo:GFP in a clone (marked by CAAX:TagBFP) simultaneously expressing *ara^RNAi^*, *caup^RNAi^*, and *mirr^RNAi^* in the lateral tissue domain at 20 hAPF. (**C**) Maps of the average tissue flow fields in control *v^RNAi^* (*N* = 8) (left) and *Tollo^RNAi^* tissues (*N* = 5) (middle) as well as the difference between *Tollo^RNAi^* and *v^RNAi^* (right) over 17:45–19:45 hAPF. Orange arrows indicate the direction and amplitude of tissue flow. Shadings indicate local indicators of spatial association (LISA) based on local Moran’s index statistic (right). Green shading denotes “High-High” (HH) spatial bins representing regions of significantly increased velocity difference; purple shading denotes “Low-Low” (LL) spatial bins of significantly decreased velocity difference. Color intensity reflects statistical significance as − log10(*p*). Red outline: ROI selected using Moran’s index *p* < 0.05. For *v^RNAi^* and *Tollo^RNAi^*-*v^RNAi^* maps are plotted onto a reference animal. Yellow dots: landmark macrochaetae. (**D**) Maps of the average isotropic deformation rate fields in control *v^RNAi^* (*N* = 8) (left) and *Tollo^RNAi^* tissues (*N* = 5) tissue (middle) as well as the difference between *Tollo^RNAi^* and *v^RNAi^* (right) over 17:45–19:45 hAPF. Grey circles indicate the amplitude of isotropic contraction, and white circles indicate the amplitude of isotropic dilation. Shadings indicate local indicators of spatial association (LISA) based on local Moran’s index statistic (right). Red shading denotes “High-High” (HH) spatial bins representing regions of significantly increased tissue dilation; blue shading denotes “Low-Low” (LL) spatial bins representing regions of significantly increased tissue contraction. Color intensity reflects statistical significance as − log10(*p*). Green outline: ROI selected using Moran’s index *p* < 0.05. Orange outline: subregion within the green ROI with contraction amplitude below −0.02 h⁻¹. For *v^RNAi^* and *Tollo^RNAi^*-*v^RNAi^* maps are plotted onto a reference animal. Yellow dots: landmark macrochaetae. (**E**) Maps of the advected average Tollo:GFP pattern (*N* = 9) at 18:45 and 19:10 hAPF. Tollo:GFP is measured at 15 hAPF and advected for a reference animal. Red outline: ROI from velocity analysis in C. Green outline: ROI from deformation analysis in D. Yellow dots: landmark macrochaetae. (**F**) Maps of the 5-MD morphodynamic domains advected onto a reference animal at 18:45 and 19:10 hAPF. Red outline: ROI from velocity analysis in C. Green outline: ROI from deformation analysis in D. Yellow dots: landmark macrochaetae. (**G**) Polar plot showing the mean tissue velocity in control *v^RNAi^* (*N* = 8) and *Tollo^RNAi^* (*N* = 5) tissues within the red ROI from C in a 2 h sliding window around the peak of Moran’s index in Figure S4A. Two-sided Mann-Whitney U test on the norm of the velocity vectors (\**p* = 4.5 × 10^−2^). (H) Bar plot of the mean isotropic contraction rate (± SEM) in control *v^RNAi^* (*N* = 8) and *Tollo^RNAi^* (*N* = 5) tissues within the green ROI from D in a 2 h sliding window around the peak of Moran’s index in Figure S4B. Two-sided Mann-Whitney U test on the isotropic component of tissue dilation (\*\**p* = 7.8 × 10^−@^). (**I**) Graph of the mean isotropic contraction (± SEM) between 16:47 and 26:47 hAPF within the green ROI from D in control *v^RNAi^* (*N* = 8) and *Tollo^RNAi^* (*N* = 5) tissues. Color shading at the top represents *p*-values under a two-sided Mann-Whitney U rank-sum test. (**J**) Graph of the mean number of extrusion events (± SEM) between 16:47 and 26:47 hAPF within the orange ROI from D in control *v^RNAi^* (*N* = 12) and *Tollo^RNAi^* (*N* = 17) tissues. Color shading at the top represents *p*-values under a two-sided Mann-Whitney U rank-sum test. (K) E-Cad:3xmKate2 and GC3Ai confocal images of a tissue region including the red ROI from C in control *v^RNAi^* and *Tollo^RNAi^* animals. Yellow dots: landmark macrochaetae. Posterior and anterior dorsocentral macrochaetae (pDC and aDC) are indicated. Scale bars: 100 µm (A, C-F), 50 µm (B, K); velocity and deformation rate are indicated (C, D).

To this end, we employed Moran’s index to quantify spatial autocorrelation.^98,99^ This approach assumes that loss of gene function generates spatially coherent differences, with adjacent domains exhibiting similar variations. By applying Moran’s index to the subtracted velocity and deformation maps (*Tollo^RNAi^* - *v^RNAi^*), we quantified the spatial autocorrelation of *Tollo^RNAi^* perturbation and identified periods manifesting the highest autocorrelated alterations in morphogenesis (Figures S4A and S4B). Velocity autocorrelation differences started with Moran’s index = 0.52 at 16:47 hAPF and increased to 0.80 at 18:43 hAPF and then remained above 0.75 for around one hour (Figure S4A). We observed low spatial autocorrelation in deformation differences at 16:47 hAPF (Moran’s index = 0.20), which increased to 0.61 at 19:11 hAPF and then plateaued (Figure S4B). These analyses therefore revealed the emergence of spatially and temporally coherent differences in both tissue flow and deformation. Interestingly, the emergence of this spatially coherent phenotype encompassed the phase of notum development during which the Tollo-enriched domain 5- MD4 undergoes coordinated tissue flows and deformations (Figures 1E and 4A; Video S1). To spatially quantify where *Tollo^RNAi^* affects tissue development, we applied local Moran’s index analysis to identify regions with statistically significant spatial clustering of differential tissue behaviors. We focused on a 2 h time interval around the maximal global Moran’s index value (18:43 hAPF for velocity and 19:11 hAPF for deformation) and defined regions of interest (ROIs) corresponding to tissue regions with significantly increased or decreased velocity (green and purple shading) and isotropic deformation rates (red and blue shading, Figures 4C, 4D and S4C and S4D; Video S3). This encompasses regions where Tollo is highly enriched within 5-MD4 and extends into 5-MD3, consistent with the graded Tollo distribution in this lateral tissue region (Figures 4E and 4F; Video S3). Quantitative analysis within these ROIs revealed two distinct aspects of the *Tollo^RNAi^* phenotype. First, *Tollo^RNAi^* caused a significant 2.9-fold increase in tissue flow velocity within 5-MD4 and the adjacent more medial 5-MD3 (red ROI, from 1.1 μm/h in control *v^RNAi^* to 3.0 μm/h in *Tollo^RNAi^*, Figures 4C, 4E-4H and S4E). The increase in tissue flow velocity is not caused by changes in cell shape or cell rearrangements, nor is it associated with changes in apoptosis rate (Figures S4F-S4I). Second, and concomitant with the larger lateral to medial flow, the tissue exhibited a 3-fold increase in isotropic contraction rate, from -0.012 h^−1^ in control *v^RNAi^* to -0.036 h^−1^ in *Tollo^RNAi^* (green ROI, Figures 4D and 4I). Interestingly, the contraction mainly occurs within 5-MD4 but extends to 5-MD5 where Tollo:GFP levels are lower from 15 to 26 hAPF (Figures 4D-4F and S4J).

To further investigate the consequences of Tollo loss of function outside the 5-MD4 domain of strong expression, we focused on regions exhibiting the highest rate of additional contraction identified by our local Moran’s index analysis (orange ROI, Figure 4D). We found that this area reduction was correlated with a 4-fold increase in cell extrusion from about 23:30 hAPF onwards (Figures 4J and S4K). Furthermore, expression of the caspase reporter GCA3i^100^ in both *Tollo^RNAi^* and *v^RNAi^* tissues established that the additional extrusions observed in this region in *Tollo^RNAi^* tissues correspond to apoptotic events (Figure 4K). Together, our analyses revealed a role for Tollo in reducing tissue flow in regions where it is highly expressed. The spatial overlap between the observed phenotypes and Tollo’s expression domain within 5-MD4 validates our computational approach to integrate morphodynamic and transcriptomic datasets. Moreover, our statistical approach also permits the objective identification of morphodynamic differences (excessive tissue contraction and apoptosis) in regions where Tollo is lowly expressed during morphogenesis.

## Discussion

Understanding the relationship between tissue morphogenesis and GRNs remains a fundamental challenge in developmental biology. To this end, we developed a comprehensive framework that links cell- and tissue-scale dynamics to spatial gene expression patterns. By combining quantitative analyses of cellular behaviors with spatial transcriptomics, we identified distinct tissue domains exhibiting coherent dynamic behaviors that are associated with specific gene regulatory programs. This was achieved through two main steps. First, we applied unsupervised spatial clustering to a large set of dynamic measurements, allowing us to segment the tissue into morphodynamic domains characterized by coherent cell behaviors. Second, we inferred GRNs associated with each morphodynamic signature using spatial transcriptomics. Supported by previous studies and additional functional validation, we confirmed that the TFs and target genes identified within these GRNs contribute to the regulation of tissue morphogenesis. Notably, our analysis uncovered a role for the TLR Tollo in preventing excessive tissue flow, tissue contraction, and apoptosis induction. By leveraging morphodynamic domains, our work provides a framework to understand how spatially organized gene expression programs coordinate diverse cellular behaviors to shape developing tissues. Because our framework integrates conserved core cellular dynamics with established GRN inference methods,^3–5,7,8,52–55,101^ we anticipate that it will be broadly generalizable. It offers both an all-in-one methodological approach to link live imaging with tissue spatial transcriptomics and a versatile platform that can be customized for user datasets.

The characterization of the Tollo loss-of-function phenotype raises important mechanistic questions regarding the relationship between gene expression patterns and tissue dynamics when inferred solely from correlation. While the role of Tollo in the lateral domain, where it is highly expressed, likely reflects local control of tissue flow, the medial tissue contraction and associated apoptosis observed in regions of relatively low Tollo expression may reflect a non-local consequence of the preceding increase in lateral flow. Our statistical framework for objectively exploring gene function therefore enables the identification of roles that could not be predicted *a priori* by correlating gene expression with local tissue dynamics alone. Such potential non-local effects may be mediated by mechanical coupling between distinct tissue regions. Interestingly, the link between spatial gene expression landscapes and inferred mechanical stresses can be probed in fixed tissues.^102^ We foresee that predicting morphogenetic phenotypes will require combining our framework with force inference methods and mechanical or agent-based modeling approaches that explicitly account for boundary conditions and mechanical effects impacting tissue dynamics at different length scales.^103–107^ Finally, the continued development of spatiotemporal transcriptomic approaches will further facilitate the characterization of bidirectional coupling between transcriptional landscapes and tissue mechanical stress across both space and time.^39–43^ In conclusion, by providing a standardized and general framework for inferring and testing GRNs based on quantitative morphometrics, our work lays the foundation for uncovering the gene regulatory principles underlying complex morphogenetic processes across diverse tissues from different species.

### Limitations of the study

To define morphodynamic domains, we have restricted our analysis to a set of broadly used kinematic and static descriptors to maintain the generality of our approach. We foresee that incorporating inferred or measured mechanical stress,^105^ along with the spatial distribution of proteins involved in the regulation of cell contractility (e.g., non-muscle Myosin II) and adhesion (e.g., cadherins), will provide additional relevant quantities for segmenting the tissue into morphodynamic domains characterized by both their kinematic behavior and mechanical properties. Fully analyzing how Tollo acts in the lateral tissue domain to regulate flow and in the more medial region to regulate contraction and apoptosis will require local inactivation of Tollo function. Within the lateral domain, clonal loss of Tollo function generates rounded clones that undergo extrusion and display ectopic Myosin II accumulation at clone boundaries (not shown). This has prevented us from performing clonal analyses to specifically assess whether Tollo acts within the lateral domain to control tissue flow. Further approaches, building on optogenetic methods to modulate Tollo function without inducing local ectopic Myosin II accumulation, will be needed to fully explore Tollo function.

## Acknowledgments

We thank François Schweisguth, the Bloomington Drosophila Stock Center, Transgenic RNAi Project at Harvard Medical School, Vienna Drosophila Resource Center, Kyoto Drosophila Stock Center, and the Developmental studies Hybridoma Bank for reagents; Hisashi Nojima and Jean-Paul Vincent for providing the Rpr and Hid reporters; Pedro Moreira Goncalves for molecular biology reagents; the Cell and Tissue Imaging Platform PICT-IBiSA, and members of the National Infrastructure France-BioImaging supported by the French National Research Agency (ANR-24-INBS-0005 FBI BIOGEN), for assistance with light microscopy and image analysis; Olivier Leroy for his contribution to the development of multichannel visualization tools for gene expression maps; Lydie Couturier for help with smiFISH; F. di Pietro for valuable comments on the manuscript. This work was supported by Institut Curie, CNRS, INSERM, ERC Advanced TIMORPH (340784) and Scaling-Sensitivity (101020243), CANCERO-INCA (PLBIO2020/BELLAICHE), ANR Labex DEEP (11-LBX-0044, ANR-10-IDEX-0001-02), Marie Sklodowska-Curie Innovative Training Network ’PolarNet’ (675407) and FRM (FDT201904008163).

## Author contributions

A.L., E.v.L., F.B., L.N., and Y.B., conceptualization, methodology; A.L., E.v.L., R.P., M.E.-C., S.R., and L.N., software; A.L., E.v.L., and L.N., formal analysis; E.v.L., M.B., L.A., F.G., J.D., and F.B., investigation; I.G., and S.P., resources; A.L., data curation; A.L., E.v.L., L.N., and Y.B., writing – original draft; A.L., E.v.L., L.A., F.B., L.N., and Y.B., writing – review & editing; A.L., E.v.L., F.B., and Y.B., visualization; F.B., L.N., and Y.B., supervision; and Y.B., project administration and funding acquisition.

## STAR METHODS

### RESOURCE AVAILIBILITY

#### Lead contact

Further information and requests for resources and reagents should be directed and will be fulfilled by the lead contact.

#### Materials availability

All unique/stable reagents generated in this study are available from the lead contact upon request.

#### Data and code availability

- All original microscopy data reported in this manuscript will be shared by the lead contact upon request.
- Any additional information required to reanalyze the data reported in this manuscript is available from the lead contact upon request.
- Code generated as part of this study will be available using a Github link upon publication.
- scRNA-seq data and vGEP profiles will be available under GEO accession number and a website, respectively, upon publication.

### EXPERIMENTAL MODEL AND SUBJECT DETAILS

#### Fly stocks and genetics

All *Drosophila melanogaster* stocks are listed in the Key resource table. Crosses were grown on standard cornmeal-agar medium at 25 °C unless otherwise specified. For temporal induction of UAS-dependent transgene expression we used the temperature sensitive *Act5C-GAL4/tub-GAL80^ts^* module.^108^ Thereto, animals were raised at 18 °C and shifted to a 29 °C restrictive temperature 48 hours prior to imaging.

### METHOD DETAILS

#### Molecular biology and transgenes

All fluorescently tagged transgenes produced for this study were generated by CRISPR/Cas9-mediated homologous recombination at their endogenous loci using the vas-Cas9 line.^109^ Guide RNAs (see Table S2) were cloned into the pCFD5 or pCFD3:U6:3gRNA vectors,^110^ while homology sequences, HR1 and HR2, were PCR amplified (see Table S2 for primers used) and cloned by SLIC^111^ into the GFP or mKate2 tagging recombination vectors, carrying a hs-mini-white cassette flanked by two LoxP recombination sites for its removal, and a N- or C-terminal fluorescent protein coding sequence.^112^ Subsequently, we removed the hs-mini-white cassette from animals using Cre-dependent recombination of the LoxP sites.^109^

To generate *UAS-ara^dsRNA^-caup^dsRNA^* and *UAS-ara^dsRNA^-caup^dsRNA^-mirr^dsRNA^*, dsRNA short hairpin sequences were selected based on sequences provided by Transgenic RNAi Project at Harvard Medical School.^113,114^ To build the double- and triple-dsRNA constructs, sequences from the multiple RNAi vector pNP^115^ including regions flanking the attB and the gypsy elements, were PCR amplified and cloned in the pCaSpeR4 vector,^116^ using pNP-mini-white primers (see Table S2). This generated a pNP-mini-white vector that enables multiple hairpin cloning and harbors a mini-white selection marker. Hairpins (see Table S2) for *ara^dsRNA^, caup^dsRNA^, mirr^dsRNA^* were sequentially cloned into this pNP-mini-white vector. The constructs were integrated at the PBac{y[+]-attP-9A}VK00027 landing site at 89E11.^117^

All constructs were confirmed by sequencing and embryo transgenesis was performed by Bestgene Inc. (Chino Hills, CA, USA).

#### Immunohistochemistry

For fixed tissue analyses of En, pupae expressing E-Cad:3xGFP were dissected 15 hours after puparium formation (hAPF), then fixed and immunostained with antibodies as described previously,^118^ using mouse anti-Engrailed (1:500) primary antibodies and goat Alexa555 anti-mouse secondary antibodies (1:200).

#### Single-molecule inexpensive fluorescence in situ hybridization (smiFISH)

For analyses of the mRNA expression of *slow, sli, pdm3, toe, crp, nw, hbs, hth, robo1, Qsox2*, *Pvr*, *dpp*, and *fj* we used smiFISH on fixed tissue expression E-Cad:3xGFP at 15 hAPF. For each gene, 25-40 probes targeting the coding sequence were designed with Oligostan^82^ (see Table S2 for probes used). All probes were designed to be compatible with a 28 nt FLAP-X secondary probe for detection, and which was added to each primary probe. This secondary probe, complementary to the FLAP-X, was flanked by two conjugated Cy3 fluorophores (Table 2). All probes were obtained in plates from Integrated DNA Technologies at 100 M in Tris-EDTA (1x TE) pH 8.0. Per gene, an equimolar mix of primary probes was prepared and diluted 1:5 in 1x TE (primary probe mix). To anneal the FLAP-X sequences of primary and secondary probes, 4 µL of primary probe mix, 1 µL of secondary probe, 2 µL of New England Biolabs Buffer 3 (10x NEBuffer 3), and 13 µL of ddH_2_0 was mixed. Annealing was performed as described previously.^82^

smiFISH was performed by adapting previously published protocols.^82^ Pupae were glued to a glass bottom MatTek dish using heptane glue, dissected in 1x PBS and fixed 20 min in 4% paraformaldehyde (PFA) in 1x PBS as previously described.^89,119^ After fixation, nota were rinsed twice 2 min with 0.1% Triton-X100 in 1x PBS. Next, nota were permeabilized by incubation for 20 min in 0.5% Triton-X100 in 1x PBS, followed by two 2 min rinses with 1x PBS. Subsequently, nota were rinsed once for 2 min, and then for 10 min in wash buffer (250 µL 8M urea, 50 µL 20x SSC, 200 µL ddH_2_O). Nota were then incubated overnight with hybridization mix (250 µL urea 8M, 50 µL 20x SSC, 125 µL Dextran Sulfate 40% w/v, 22 µL Vanadyl complex, 9.4 µL competitor DNA (1:1 of 10 mg/mL sheared salmon sperm DNA:10 mg/mL *E. coli* tRNA), 15.6 µL smiFISH annealed primary & secondary probe, 28 µL ddH20) at 37°C in an opaque humid chamber. Following hybridization, nota were rinsed for 2 min in wash buffer and subsequently washed three times 10 min in 2x SSC. Finally, nota were rinsed 2 min and incubated for 10 min in 0.1% Triton-X100 in 1x PBS, after which nota were mounted in Vectashield mounting medium without DAPI.

#### Image acquisition

Microscopes used for imaging included an inverted spinning disk wide borealis confocal microscope CSU-W1 (Andor/Roper/Nikon) with sCMOS camera (Orca Flash4, Hamamatsu), an inverted spinning disk wide confocal microscope CSU-W1 (Roper/Nikon/GATACA) with sCMOS camera (BSI camera with 95%QE), and an inverted spinning disk wide homogenizer confocal microscope CSU-W1 (Roper/Zeiss/Spark) with sCMOS camera (Orca Flash4, Hamamatsu) using either a 40x/1.3 OIL DIC H/N2 PL FLUOR or 40x/1.4 OIL DIC H/N2 PL FLUOR objective. For live imaging of notum tissue dynamics, pupae expressing E-Cad:3xGFP, E-Cad3xmKate2, or co-expressing E-Cad:3xmKate2 and the caspase sensor GC3Ai were staged and mounted on a glass coverslip as described in.^120,121^ Pupae were typically imaged for 15-20 hours starting around 15 hAPF by multi-position time-lapse microscopy. For each position Z-stack (comprising 30–50 optical slices with a 0.5 µm step size) time-lapse images were acquired every 5 minutes at 0.161 µm/pixel resolution. Metamorph autofocus detection was used to keep the focus of the tissue for the duration of the imaging. For the compilation of the notum gene expression atlas as well as for validation of predicted gene expression profiles, multi-position, Z-stack images of fixed or live tissue were acquired at 15 hAPF as previously described.^6^

### QUANTIFICATION AND STATISTICAL ANALYSIS

#### Image pre-processing (projection, stitching and quantification)

Time-lapse and protein expression atlas image stacks were projected using a custom MATLAB code, which detects the apical E-Cad:3xmKate2 or E-Cad:3xGFP signal. The resulting images were stitched in Fiji,^122^ after which the background was subtracted using rolling ball subtraction (radius 50 pixels). The brightness/contrast intensities were adjusted (auto) and original 16-bit images were converted to 8-bit. For smiFISH images, the signal was segmented by first applying a Laplacian of a Gaussian filter in Fiji (Mexican Hat Filter, kernel size 1), and local maxima were identified using the Find Maxima plugin (Prominence parameter value: 3 for *robo1, hbs, slow*, *pdm3*, 10 for *dpp,* 40 for *toe*). Images of highly expressed genes (*nw, fj, Qsox2, sli, hth, crp, Pvr*) were processed following to the same procedure as the fluorescent protein images.

##### Velocity field quantifications

Particle image velocimetry (PIV) was performed to generate tissue-wide velocity fields in an Eulerian reference frame as previously described.^123^ The analysis utilized a multi-pass algorithm implemented in a custom MATLAB script, with a 50% overlap between adjacent interrogation windows to decrease variability. An initial pass with a large 128×128 pixel (0.161 µm per pixel) interrogation window was used to robustly capture large-scale displacements. This was followed by a refinement pass using a smaller 64×64 pixel window to achieve higher-resolution velocity measurements. The resulting discrete velocity vectors for each animal were first interpolated using a spline-based method to generate a continuous velocity field, v*_a_*(x, *t*)(where x and t denote the spatial coordinates and developmental time, respectively, and *a* denotes the animal on which the quantification was performed), across the tissue. These individual velocity fields were then averaged across all animals to create an ensemble average of tissue flow dynamics. To characterize the overall temporal evolution of morphogenesis, the magnitude of the averaged velocity vectors was spatially averaged across the entire notum at each time point, using a 4-hour sliding window.

##### Time registration

To enable comparison and averaging across different animals and experimental conditions, all time-lapse movies were registered in time. Two distinct registration methods were employed, chosen based on the duration of the recordings and the specific analysis being performed. The primary synchronization method was based on tissue flow dynamics (see section Velocity field quantifications) and was used for analyzing tissue-scale morphogenesis. For each of these movies, the mean posteroanterior tissue velocity was calculated. The reference time was defined as the time point at which the velocity reached three-quarters of its peak value and was set to 22 hAPF. A secondary method was required for the cell extrusion analysis that was performed on shorter time lapse movies and did not always capture the full posteroanterior tissue velocity profile. Time registration was performed using the stereotyped sensory organ precursor (SOP) division events as a temporal landmark. The synchronized set of movies, aligned using tissue flow velocity, served as a reference to establish a high-confidence timeline of SOP divisions. This analysis revealed that temporal registration of movies using the third division of the third row of SOPs resulted in the lowest overall timing variability. The median time of this division across samples was determined to be 17:30 hAPF. For each movie to be registered, the frame corresponding to this reference SOP division was manually identified. This process anchored all movies to a common reference time point and ensured that all time-lapse movies were accurately aligned temporally, allowing for the averaging and comparison of morphogenetic dynamics across all animals.

##### Space registration

To correct for variations in tissue size and orientation, all movies and images were aligned to a common spatial coordinate system. This was achieved through a two-step process using a suite of custom MATLAB scripts.^4^ First, a canonical reference "archetype" was generated to define a standardized tissue geometry. Second, all individual images and movies were spatially registered to this archetype via an anisotropic scaling transformation.

The generation of the archetype began with the manual identification of four to eight invariant macrochaetae and two points defining the tissue’s midline, the axis of symmetry (see Figure 1B). These landmarks were manually identified using a custom graphical user interface on 270 E-Cad:3xmKate2 and E-Cad:3xGFP images at 15 hAPF. Landmark coordinates from tissue images were centered by translating the origin to the centroid of the macrochaetae. An initial, provisional archetype was computed by averaging these centered landmark positions. Each tissue’s landmarks were then homothetically rescaled to best fit this provisional archetype via a least-squares minimization. The final, canonical archetype was defined as the average of these rescaled landmark positions:

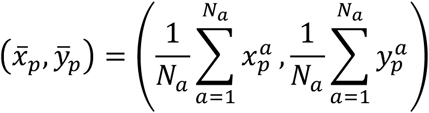

where *p* = 1, …, *N* indexes the landmarks (up to 8 macrochaetae), *a* = 1, …, *N_a_* indexes the animals used for archetype generation, and (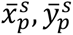) denotes the archetype coordinates for landmark *p*.

For the subsequent spatial registration of each individual tissue and movie, landmarks were identified in the same manner and aligned to the canonical archetype. For movies, animals were rescaled at 17:40 hAPF (see section Grid generation and temporal averaging). After converting landmark coordinates to centered microns, specific scale factors for the x- and y-directions were computed by finding a multiplicative factor to minimize the dispersion of macrochaetae of *a* from the archetype:

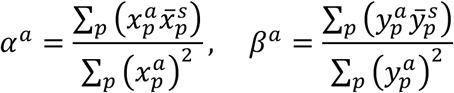

where(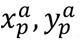) are the coordinates of landmark *p* in animal *a*, and (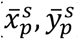) are the corresponding archetype coordinates. This transformation anisotropically stretches or shrinks the tissue’s spatial coordinates to align its landmarks with the archetype.

##### Image segmentation and cell tracking

Cell segmentation and tracking were performed using a custom computational pipeline to extract E-Cad:3xGFP cell outlines and lineages from the time-lapse movies. First, individual cells were segmented in each frame using a U-Net-based model built upon the Cellpose framework.^124^ To achieve high accuracy on our specific dataset, the pre-trained ’cyto2’ model was fine-tuned using a human-in-the-loop correction strategy, where manual corrections were iteratively used to retrain the model using a custom code. The resulting segmentation masks were then skeletonized to generate precise, single-pixel-wide cell outlines. Following segmentation, cell tracking was performed to reconstruct cell lineage trees and trajectories over time. This was achieved using a custom C++ pipeline implementing a tracking algorithm described previously.^4^

##### Cell-based quantification of morphodynamics

The quantitative analysis of morphogenesis was based on the tensor decomposition formalism described previously.^4^ This framework additively decomposes the total tissue deformation rate tensor (G) obtained from segmentation and tracking into the sum of the deformation rates associated with each cell process: cell rearrangements (R), cell shape changes (S), cell divisions (D), and cell extrusions/apoptosis (A) as well as non-biological parameters extracted from tracking and segmentation artefacts: cells appearing in the tracking (N), fusions of neighboring cells (F), and flux of cells in and out of the field (J), which are negligible in the Lagrangian frame of reference:

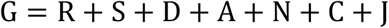

To facilitate statistical analysis, which requires scalar inputs, a set of scalar features from the vector- and tensor-based morphogenetic data were calculated. From the velocity vector v = (v_x_, v_y_): its magnitude, or speed, as 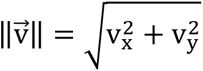. Rates of discrete events like cell apoptosis k_A_ (h^-1^) and divisions k_D_(h^-1^) were normalized by the number of cells present at the beginning of the measurement window (typically 4 hours). From each 2×2 deformation tensor T, we extracted two key scalar metrics: Tr[T] (h^-1^), representing the isotropic component of deformation rate (area change) for a tensor T, and Dev[T] (h^-1^), representing the deviatoric component of deformation rate for a tensor T (local elongation/contraction). These two scalar metrics are indicated as T → dilation for Tr[T] and T → elongation for Dev[T]. This process yielded a set of 15 primary scalar properties. These included T → dilation for Tr[T] and T → elongation for: the total deformation tensor G as well as for each of its constituent cellular behaviors: R, S, D, A. The remaining five properties were the magnitude of cell velocity vectors 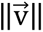, normalized division rate k_D_ (h^-1^), normalized apoptosis rate k_A_ (h^-1^), mean apical cell area (µm^2^), and cell shape anisotropy calculated as 1 − (minor axis/major axis) (unitless) (see Table S1 for quantitative descriptors used).

##### Grid generation and temporal averaging

All quantified data were subjected to spatiotemporal averaging. First a grid was initially overlaid on the live-imaging data at 17:40 hAPF to comprise all movies. The same grid was used on the gene expression atlases and smiFISH images, as tissue flow and deformation from 15 hAFP to 18 hAPF are minimal.^4^ Unless mentioned otherwise, all subsequent calculations were performed in a Lagrangian reference frame, meaning the grid follows cell movement obtained using cell tracking from the time-lapse movies. Spatially, quantities were averaged within Lagrangian bins of 40 µm x 40 µm, with a 50% overlap between adjacent bins.^4^ Temporally, a sliding window average was applied. The window size was set to 4 hours for the broad morphogenetic analysis (Figures 1 and S1; Video S1 and S2) and to a shorter 2-hour window for the tissue velocity, deformation and delamination analyses (Figures 4 and S4; Video S3). As a final quality control step for each animal, a weight for each bin representing the fraction of the bin occupied by cells, see Guirao et al.^4^ was calculated for each bin. Bins with a low weight (< 0.3 for movies, <0.9 for images) were excluded from the analysis for each animal.

##### Tissue based quantifications of morphogenesis

We computed the tissue flow velocity for each spatial bin using the defined grid structure and utilized the spatial gradients across this grid to determine a segmentation free deformation rate. The tissue flow velocity field was used to extract the deformation rate tensor (ɛ) for each animal from its respective continuous velocity field. This was calculated from the velocity gradient tensor, L = ∇v. The deformation rate tensor ɛ is the symmetric part of 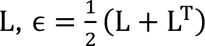. The trace of the averaged tensor Tr[ɛ] represents the isotropic rate of area change (local tissue contraction or expansion), while its deviatoric (traceless) part describes the anisotropic component of deformation (shear and elongation). For visualization, both velocity vectors v and deformation rate tensors ɛ were averaged across all time-lapse movies of a given genotype. These fields were visualized using the AnimalProcessing package using Napari^125^ available as part of this publication. To highlight coherent patterns of flow and deformation, a local consistency filter was applied based on inter-replicate variability. The consistency was determined at each time point for each spatial coordinate (bin) using the local mean (μ) and standard deviation (σ) of the measurement calculated across all time-lapse movies of a given genotype. Velocity vectors are highlighted in grey if their average magnitude exceeded 0.5 σ.

##### Averaging quantities across animals

Following temporal and spatial registration, the filtered spatiotemporal fields for all quantified properties were averaged across animals within each experimental condition. This procedure created a "mean animal" by computing an average weighted by the weight of each bin, thus ensuring that regions with more cellular data contributed more significantly. Once the mean animal was calculated, a final filtering step was applied: patches with a weight below 0.15 and any remaining isolated patches were excluded.

For the generation of the velocity and deformation fields visualization in the Eulerian frame of reference (Figures 1D and Video S1), patches with a weight below 0.55 were excluded. Manually defined circular masks were applied to remove regions to enhance visualization (e.g., half a row in the midline).

For the generation of the tensor decompositions visualization in the Lagrangian frame of reference (Figures 1E and Video S2), patches with a weight below 0.15 were excluded. Values above 0.98 percentile were clamped to enhance visualization. Origins not overlapping with the cells of the reference animals were removed due to the impossibility to advect them by tissue deformation.

Separately, the average protein expression atlases and smiFISH images were generated by averaging the profiles across a minimum of three replicates (*N_a_* ≥ 3). After averaging, these static atlas intensities were rescaled using percentile normalization (0.05-0.995).

##### Dimensionality reduction and identification of morphodynamic domains

Principal component analysis (PCA) was based on a set of 15 quantitative morphogenetic properties extracted for each spatial bin (40 µm x 40 µm). Each of these 15 properties was measured at two distinct time windows, an "early" phase (18-22 hAPF) and a "late" phase (22-26 hAPF), creating a final feature vector of 30 variables per spatial bin. Prior to PCA, the 30 features were standardized across all spatial bins using the ‘StandardScaler‘ from the scikit-learn library to ensure each feature had a mean of 0 and a standard deviation of 1. PCA was then performed on this scaled feature matrix using scikit-learn. The contribution of each original feature (loadings) was compared to interpret their biological significance and the component scores were visualized as spatial maps. For subsequent clustering analysis, the top 14 principal components were retained, which collectively explained over 90% of the total variance in the data.

To identify spatially contiguous morphodynamic domains, we performed spatially constrained agglomerative hierarchical clustering on the top 14 principal components derived from the morphometric data. First, a spatial connectivity graph was constructed using the radius_neighbors_graph function from scikit-learn, with a radius set to √2. This constraint ensures that the clustering algorithm can only merge bins that are spatially adjacent (queen contiguity), thereby enforcing the creation of unfragmented domains. The AgglomerativeClustering function from scikit-learn was then used with Ward linkage, supplied with the precomputed connectivity graph to cluster bins based on spatial proximity.

To objectively determine the optimal number of clusters (morphodynamic domains), clustering was performed for a range of 2 to 13 partitions. For each partition, the silhouette score from scikit-learn was computed. The clustering hierarchy was visualized using a dendrogram (see Figures S1E and S1F).

To enhance the spatial coherence of the resulting domains, a custom iterative post-processing algorithm was applied to the cluster assignments for each partition. This algorithm refines the cluster map by eliminating small, isolated patches (rook contiguity). In each iteration, a spatial bin is identified as isolated if none of its four cardinal neighbors (up, down, left, right) share its cluster label. Such isolated bins are then reassigned to the majority cluster label of their four cardinal neighbors. In the case of a tie, the neighborhood is expanded to include the four diagonal neighbors (8-connectivity), and the bin is reassigned to the majority cluster of this expanded 8-neighbor set, provided a clear majority exists. This process was repeated for a maximum of 5 iterations or until the cluster assignments stabilized. After this procedure, cluster labels for each partitioning were reindexed to form a consecutive sequence starting from 1. The final spatial organization of the resulting domains was visualized as colored maps overlaid on the tissue outline.

#### Single cell sequencing

Single-cell RNA sequencing was performed on five replicates (four biological replicates, with replicate 3 and 4 being technical replicates of the same biological sample). For each biological replicate 40–45 female *w*^1118^ pupae were dissected at 15 hAPF in ice cold 1x PBS without CaCl_2_ and MgCl_2_ (DPBS). Nota were dissected as previously described.^118^ After dissection nota were transferred to a LoBind Eppendorf tube and dissociated in 150 μL TrypLE solution at 29 °C for 15 min. The tube was flicked, just prior to incubation at 29 °C as well as after approximately 7 min of incubation. Following dissociation, the cells were vortexed at 25 Hz for 30 s and the enzymatic reaction was quenched by the addition of 750 μL of freshly prepared 2% bovine serum albumin (BSA) in 1x DPBS. Subsequently, cells were pelleted by centrifugation at 400 rcf for 5 min at 4 °C. The 30 μL cell pellet was then resuspended in 100 μL of freshly diluted 0.5% BSA in 1x DPBS and vortexed at 25 Hz for 30 s. To assess cell viability and the cell count, cells were diluted 1:1 in a 0.4% Trypan Blue solution, added to a hemocytometer and inspected using an upright widefield Leica microscope with 20x/0.4 HCX PL FLUOTAR CORR (506242) objective. Typically, this yielded a cell suspension of 250-500 cells/μL, with ∼85% single cells and ∼95% viability.

Single-cell libraries for sequencing were prepared using the 10x Genomics Chromium platform with Chromium 3’ v3 chemistry. Targeted cell counts were 10,000 (biological replicates 1 and 2), 20,000 (biological replicate 3 and its technical replicate, 4), and 2,000 (biological replicate 5). The resulting cDNA was amplified for 12 cycles, and libraries were sequenced on a NovaSeq 6000 platform (Illumina) in PE 28-8-91 with a coverage of 40,000-60,000 reads/cell. scRNA-seq data were deposited in NCBI GEO with accession number GSE328433.

#### scRNA-seq data processing and analysis

Raw scRNA-seq data were processed using the Salmon Alevin module^126^ with the chromium V3 flag, corresponding to libraries prepared using Chromium 3’ v3 chemistry. Reads were quantified against a decoy-aware Salmon index built from the *Drosophila melanogaster* transcriptome (v6.22). The index, which also included genomic decoy sequences to reduce spurious mappings, was generated using a k-mer size of 31. The resulting data for each replicate was subsequently imported into R using the Bioconductor tximport package (1.26.1)^127^ and gene identifiers were converted to unique gene symbols with the Bioconductor org.Dm.eg.db annotation package (3.16.0) (Bioconductor - org.Dm.eg.db). This resulted in five individual datasets, for which we created initial objects using the Seurat package,^81^ retaining genes expressed in at least 5 cells and cells with at least 200 detected features.

Next, we applied a multi-step quality control pipeline to each of the five datasets independently. First, cells were filtered based on their Unique Molecular Identifier (UMI) counts and mitochondrial DNA content. To account for batch effects and technical variability across replicates, UMI count thresholds, were tailored for each sample: 12,000-110,000 (replicate 1), 14,000-120,000 (replicate 2), 7,500-85,000 (replicate 3), 7,500-100,000 (replicate 4), and 12,000-85,000 (replicate 5). A maximum mitochondrial content of 10% was uniformly applied for all samples. We then removed potential doublets using the DoubletFinder package (2.0.3)^128^ within each dataset. Finally, we excluded cells with low-complexity libraries, defined as having more than 12.5% of UMIs originating from a single gene. After applying these filters, the five datasets were merged. Our scRNA-seq provided 12,187 high quality cells and 9,699 genes. To mitigate batch effects and technical variability originating from the different sequencing runs, we integrated the five filtered datasets using the anchor-based workflow in Seurat v3.^81^

For each replicate independently, gene expression counts were log-normalized (NormalizeData, LogNormalize method) using the median UMI count of the replicate as a scaling factor. The top 2,000 highly variable genes (HVGs) were identified using variance-stabilizing transformation (FindVariableFeatures, vst method), and expression values were scaled across all genes (ScaleData). We then performed principal component analysis (PCA) on the scaled data and used the top 30 principal components (PCs) to identify integration anchors across the five replicates. Ultimately. We then performed clustering using the Leiden algorithm (FindNeighbors followed by FindClusters with a resolution of 0.2). For visualization, we used UMAP.^129^ We assigned cell type identities to the resulting clusters by manually inspecting the expression of a panel of canonical marker genes for known cell types associated with the pupal notum epithelium. Notum epithelial cells (7,610 cells) were identified by the expression of: *hh*, *pnr*, *caup*, *eyg*, *ush*, and *mirr*. Non-notum epithelial cell types, which were subsequently excluded from downstream analysis, included: neck epithelial cells (434 cells, *Dfd*, *oc*), myoblasts (*twi*, *sns*, *kon*, *lmd*, *Him*) and tracheal cells (*htl*) were clustered together (2,561 cells), neuronal sensory organ precursors (174 cells, *sens*, *mira*, *ase*), and a remainder of unassigned cells (1,408 cells) was left unannotated (Figures 2A and S2A).

#### Spatial transcriptome reconstruction

##### Spatial reference atlas construction

To build the spatial expression reference atlas for our spatial transcriptome reconstruction, we generated at least 3 images for *al*, *ara*, *B-H1*, *bi*, *Diap1*, *esg*, *exd*, *eyg*, *h*, *lgs*, *mirr*, *Pax*, *ptc*, *salm*, *sd*, *rpr*, *ush*, *wg*, *sr*, *tsh*, *hh*, *usp*, *hid*, *en*. Images were spatially registered and rescaled using macrochaetae and averaged as described in the Space registration section. These final, registered, averaged, and rescaled atlas images were then discretized into common 40 µm x 40 µm spatial bins, identical to the grid of the common archetype hemi-notum used for morphodynamic measurements.

##### Mapping single cells

A custom computational pipeline was developed to reconstruct a genome-wide spatial map of gene expression in the hemi-notum by projecting the 7,610 high-quality epithelial cells from the scRNA-seq dataset onto the spatial bins of the reference atlas. To obtain reliable patterns, a more stringent gene-level filter was applied, retaining only genes expressed in at least 50 cells and maximal expression at least above 10 UMIs in one cell which yielded 4,312 genes. These cells were then mapped onto the tissue using the following algorithm:

1. *Single-cell data preprocessing:* The raw single-cell count matrix (*X*) for the 7,610 cells by 4,312 genes was first normalized for sequencing depth by dividing each cell’s counts by its total UMI count, then multiplying by a scale factor (10,000) to create a scaled expression matrix (*X^scaled^*). The subset of genes common to both the atlas and the single-cell data (*G_T_*) was identified. For these training genes, the corresponding expression data was also log-transformed 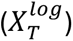.
2. *Correlation-based mapping:* The similarity between each spatial bin and each cell was calculated by computing Pearson correlation matrix (*R*) (461 bins x 7,610 cells) between the log-transformed expression profile of the training genes in each cell i (*X*_567,*T*_) and the corresponding gene signature in each spatial bin k of the atlas (*A_T_*_,*k*_):

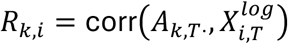

where *k* is the spatial bin index, and i a single cell. To refine the mapping, this matrix was then filtered. We then calculated a quantile threshold (*q*) from its distribution of correlation values based on the validation set (see next section). Only positive correlation values above this threshold were retained, creating a mapping matrix (*R^map^*).
3. *Weighted reconstruction:* The full transcriptome for each spatial bin (*P_st_*) (461×4,312 positions by genes) was computationally reconstructed by performing a weighted average of the normalized expression profiles (*X_scaled_*) of all single cells, using the filtered correlation scores from *R_map_* as weights:

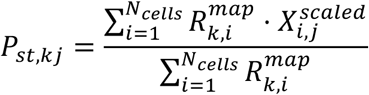

Using this approach, we reconstructed the virtual gene expression patterns (vGEPs) for 4,312 genes expressed in the notum epithelial cell population at 15 hAPF, based on 3,765 mapped cells, while 3,845 cells were discarded due to low correlated with each spatial bin.

##### vGEP validation and parameter optimization

The accuracy of the reconstructed vGEPs was validated by comparing them against the independent ground-truth validation set of 13 smiFISH RNA expression profiles (*dpp, Pvr, crp*, *fj*, *hbs*, *hth*, *nw*, *pdm3*, *Qsox2*, *robo1*, *sli*, *slow* and *toe*) as well as 7 protein pattern profiles (*C15, caup, Dl, WntC, vvl, 18w, and svp*), that were not included in the spatial expression reference atlas. At a later stage, the Tollo expression pattern was added for comparison to vGEP, without including it in the validation.

This validation set was used to quantitatively optimize the parameters of our reconstruction pipeline. We optimized key upstream parameters, including those defining the epithelial scRNA-seq subset (e.g., number of PCs, HVG selection methods, inclusion of the neck population and unassigned cluster in the potentially mapped cells), as well as the correlation-based mapping threshold (*q*), which determines the final number of cells mapped to each position. The final parameters were chosen by selecting the combination that maximized the composite score (mean + median) of Pearson correlation coefficients between the reconstructed vGEPs and the experimental validation set.

To evaluate the performance of our reconstruction pipeline, we benchmarked it against two spatial reconstruction algorithms: NovoSpaRc^20,83^ and Perler^21^. Both NovoSpaRc and Perler methods were systematically optimized to achieve their best possible performance on our dataset using the same validation set and optimization metric for a fair comparison. For NovoSpaRc, we performed an extensive grid search over its key hyperparameters. The search space included the alpha parameter (20 values from 0.0 to 1.0), which balances the atlas-based and transcriptomic similarity costs; the entropy regularization term, epsilon (3 log-spaced values); the number of spatial neighbors (5, 10, 15, 20, 25); and the number of transcriptomic neighbors (5, 8, 10, 12, 24). We also tested two different input representations of the single-cell data: the log-normalized expression matrix and its principal components. The optimal combination of parameters was selected based on the Pearson correlation coefficients between the reconstructed patterns and the experimental expression patterns of the validation set. For Perler, we performed a similar optimization by conducting a grid search over its primary hyperparameter, the number of metagenes, testing all integer values from 5 to 22. The optimal number of metagenes (*n* = 7) was chosen using the same evaluation metric of sum of mean and median of Pearson correlations against the validation set.

#### Analysis of the transcriptional landscape associated with morphodynamic domains and predicting GRNs associated with tissue flow and deformation

##### Identification of domain enriched genes

To identify genes specifically expressed in epithelial morphogenetic domains, we employed the Tau (τ) tissue specificity index.^85^ Critically, the τ score is highly sensitive to the composition of the reference samples (clusters) included in its calculation. Initial analysis restricted solely to the five epithelial domains was susceptible to retaining non-specific or weakly expressed markers (e.g., the neck marker *Dfd*), thereby introducing artifacts. To rigorously control for non-specific expression and ensure the fidelity of the domain-specific gene set, we constructed an expanded unified reference expression matrix that included non-epithelial cell type clusters, thereby removing the influence of background expression profiles.

This matrix integrated the average expression profiles of the 5-MD spatial morphogenetic domains, a spatial "background" cluster aggregating all transcriptomic bins falling outside these 5 domains, and scRNA-seq pseudobulk profiles for distinct cell types (muscle and trachea, sensory organ precursors, epithelium neck, and unassigned cells) obtained by averaging scaled cell expression belonging to each cell type. The later clusters, used as references, were excluded from the heatmap representation to focus on morphodynamic domains of interest. Prior to pseudobulk aggregation, scRNA-seq raw counts were normalized to a target sum of 10,000 counts per cell using Scanpy,^130^ for the same 4,312 genes in the spatial transcriptome, consistent with the UMI count normalization applied to each spatial transcriptome bin. Using the tspex package, we then computed τ ^86^for all spatially reconstructed genes based on their expression across the spatial domains. To strictly exclude genes primarily expressed in non-epithelial cells or surrounding background, we applied a "maximum expression" filter, retaining only genes (992 genes) whose maximum expression value across the entire unified reference matrix occurred specifically within one of the 5-MD morphodynamic domains. This step ensured that genes highly expressed in the neck, muscles, trachea, or the "outside" spatial cluster were removed from the domain-specific analysis.

For the retained genes, we analyzed the distribution of τ scores, which revealed a bimodal profile distinguishing broad from specific expression. We applied Otsu’s thresholding method to this distribution to objectively define a specificity cutoff (τ > 0.36). 259 genes exceeding this threshold were classified as domain specific. Finally, we intersected this list with a comprehensive database of *Drosophila* transcription factors (TFs),^131^ resulting in 38 domain-specific TFs serving as seeds for gene regulatory network (GRN) inference. Each TF was assigned to the morphogenetic domain where its expression was maximal. For visualization of expression patterns and pseudobulk expression, values were log-transformed (log1p) and standardized (Z-score) across all conditions, and displayed as a heatmap.

##### Inference of spatially resolved GRNs

To reconstruct the regulatory architecture governing each morphogenetic domain, we integrated the spatial specificity results with a consensus GRN inference. Unlike the τ score, which is optimized to identify genes with high, spatially restricted expression, the subsequent identification of regulatory targets (TF-target links) must account for both activation (positive co-expression) and repression (negative co-expression). Therefore, we did not impose a τ specificity filter on the target genes to allow for the comprehensive detection of regulatory links, including those involving transcriptional repression. We used the GENIE3 algorithm,^86^ implemented in pySCENIC,^132^ to predict TF-target regulatory links based on non-linear co-expression patterns in the scRNA-seq data. We aggregated results from 101 independent GENIE3 runs. A regulatory link between a TF and a putative target was retained only if it was present in at least 80% of the independent runs. For each of the 38 domain-specific TFs identified in the spatial analysis, we retrieved their putative targets and selected the top 10 ranked by their mean feature importance score. This yielded a meta-network of 38 TFs and 598 unique target genes. We subsequently applied Leiden community detection to this network to identify modules of co-regulated genes associated with specific morphogenetic behaviors. Community detection was performed using the Leiden algorithm^133^ with a resolution parameter of 1.0.

##### Quantification of tissue dynamics in v^RNAi^ and Tollo^RNAi^ conditions

The analysis of the *Tollo^RNAi^* phenotype was performed in a Eulerian reference frame to better follow the effect of Tollo downregulation on live images. We followed a multi-step procedure to identify and quantify significant perturbations on a primary batch of 8 control (*v^RNAi^*) and 5 *Tollo^RNAi^* animals. We first computed the average velocity and deformation maps for the *Tollo^RNAi^* and *v^RNAi^* time-lapses movies. Differential maps were then generated by subtracting the *v^RNAi^* average from the *Tollo^RNAi^* average at each corresponding time point and spatial bin. To distinguish spatially homogeneous phenotypic regions from random developmental noise, we employed Global Moran’s index, a statistical measure of spatial autocorrelation.^98^ The Global Moran’s index is calculated as:

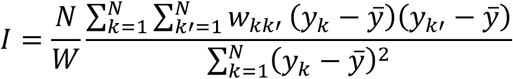

where *y_k_* is the value in a given spatial bin *k*; *ȳ* is the mean value across all bins; *N* is the total number of bins; *w_kk_*_′_ is the spatial weight between bin *k* and *k*′ constructed based on queen contiguity (where each bin of 40 µm x 40 µm is connected to its eight adjacent and diagonal neighbors) and *w* is the sum of all weights (∑*_k_*_,*k*′_ *W_kk_*_′_).

Global Moran’s index was calculated for each time point of the different movies (which were generated using a 2-hour sliding window). The analysis was conducted on two scalar fields derived from the differential tensor maps: the magnitude of the velocity vector 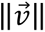 and the isotropic component of the deformation rate tensor Tr[ε], yielding a temporal profile of spatial autocorrelation for both quantities. We used the maximal value of Moran’s index (peak of global spatial autocorrelation) to select the time window at which to study the knockdown-induced perturbations (Magnitude of the Velocity: 17:45-19:45 hAPF with Moran’s index equals to 0.8, isotropic Deformation rate 18:10-20:10 hAPF with Moran’s index equals to 0.6).

To pinpoint the specific locations of the coherent phenotype within the differential maps, we applied local indicators of spatial association (LISA), using the local Moran’s index statistic for each spatial bin *k*.^99^ It is defined as:

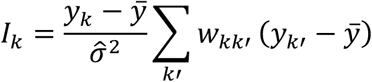

where 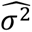 is the estimated variance of *y* and other terms are as defined previously.^99^

LISA identifies statistically significant spatial clusters by assessing the value of each bin in the context of its neighbors. We classified each bin into one of four categories: "high-high" (a high value surrounded by high values), "low-low" (a low value surrounded by low values), "high-low," and "low-high." The statistical significance of these clusters was determined by a permutation test (n = 10^5^ permutations, PySAL package esda2.5.1). We defined regions of interest (ROIs) for subsequent targeted quantitative comparisons of the magnitude of the velocity vector and the isotropic component of the deformation rate tensor as spatially contiguous regions of "high-high" or "low-low" bins with a *p*-value < 0.05 (Red and green ROIs, in Figures 4C and 4D, respectively). Within these regions of interest, we looked at the average velocity and the average isotropic dilation. We compared average tissue dilation and velocity amplitude across conditions using a two-sided Mann-Whitney U test.

#### Quantification of cell extrusion

Cell extrusion was quantified within the “high-high” ROI for the isotropic component of the deformation rate tensor. Within this region, we selected values of isotropic dilation below 0.02 (h^-1^, orange ROI in Figure 4D) and we defined an "advected region" that followed cells within the Moran ROI region as the tissue moved in time using tracking. The dataset was expanded to include a total of 12 *v^RNAi^* and 17 *Tollo^RNAi^* movies recorded in the region of increased contraction in *Tollo^RNAi^* animals. The additional animals were spatially and temporally registered as described in the space and time registration sections. Extrusion events were then counted manually. Extrusion rate was calculated by counting extrusion events in each frame (5 min interval), averaged over the 2-hour time window, and normalized by the total number of cells in the ROI. The rates were averaged across each animal and compared using a two-sided Mann-Whitney U test at each 2-hour period.

#### Statistics, figure, and video display

Statistical tests used to assess significance are indicated, and the number of samples (*n*) or animals (*N*) is reported in each figure legend and in the Quantification and statistical analysis section. Data are shown as mean ± SEM or mean ± SD. Confocal images and videos associated with this work reflect maximum intensity projections of z-stacks that were subjected to processing (denoising, contrast enhancement) in Fiji, PaintShop Pro or Photoshop for display purposes. Data plotted in Figures 1D, 4C and 4D, as well as Videos S1 and are acquired from analyses obtained from an Eulerian reference grid. Data plotted in Figure 1E, Video S2 are acquired from analysis obtained from a Lagrangian reference grid. Outlier grid positions (e.g., at the tissue midline) were manually removed to enhance visualization.

## Key resources table

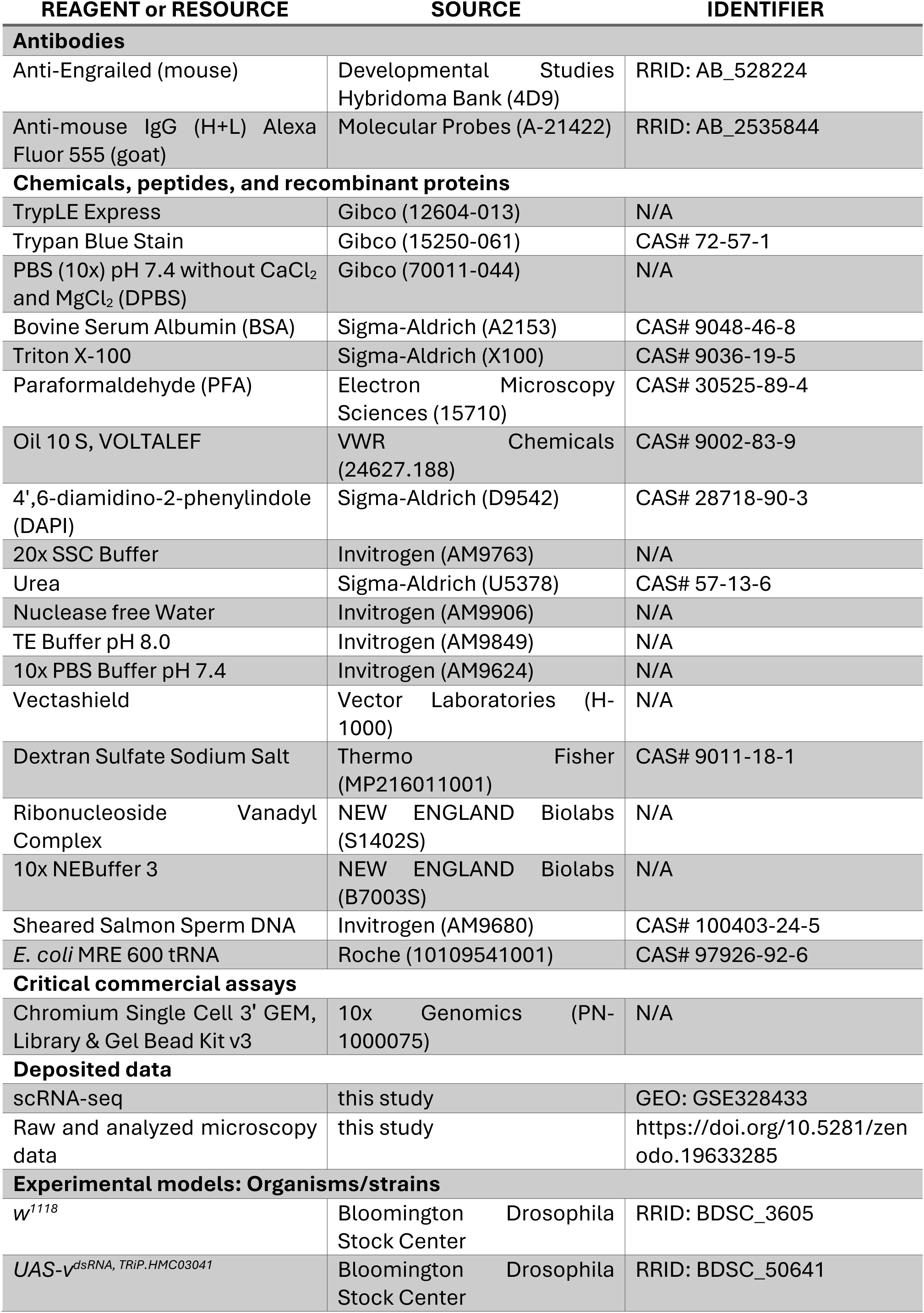
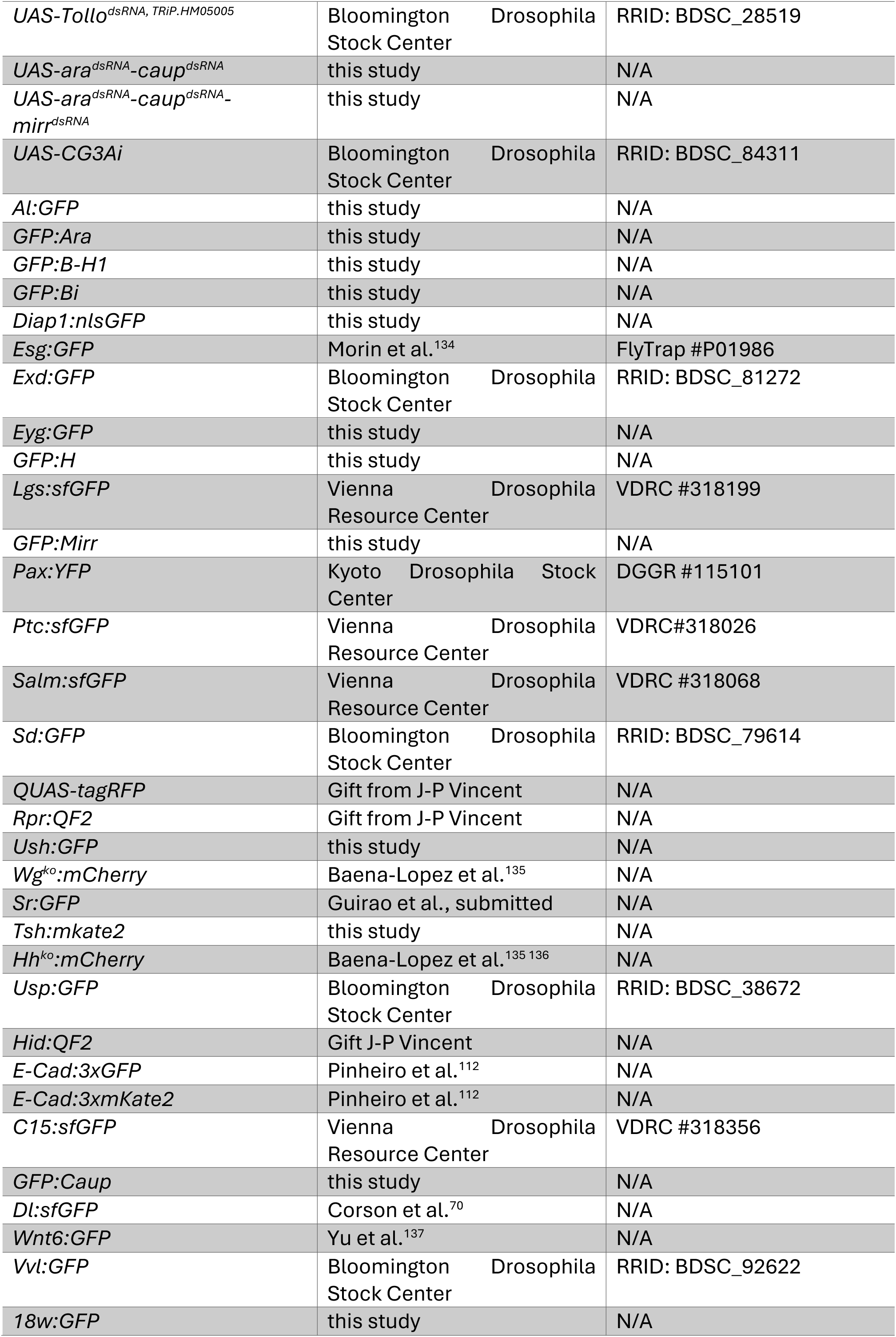
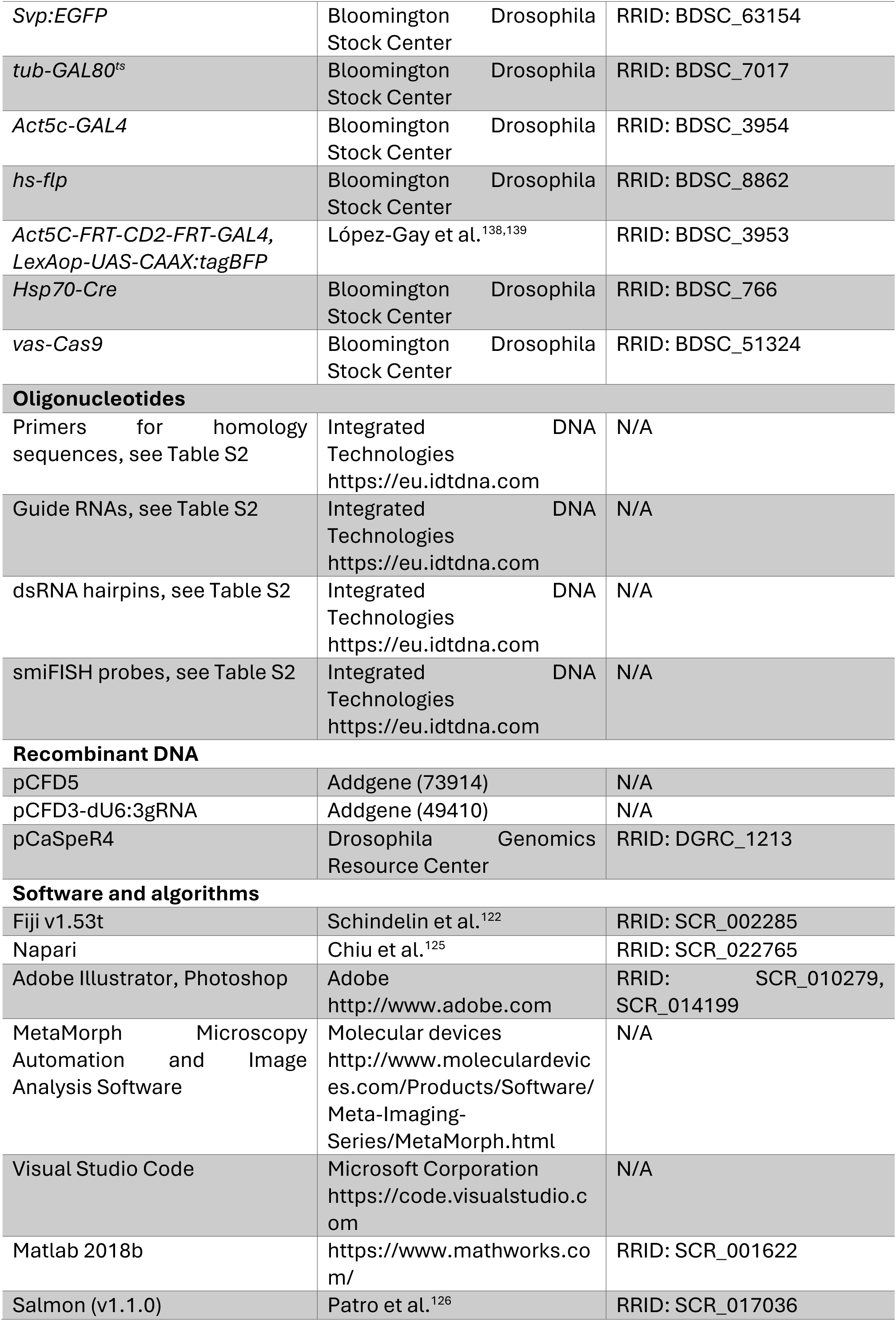
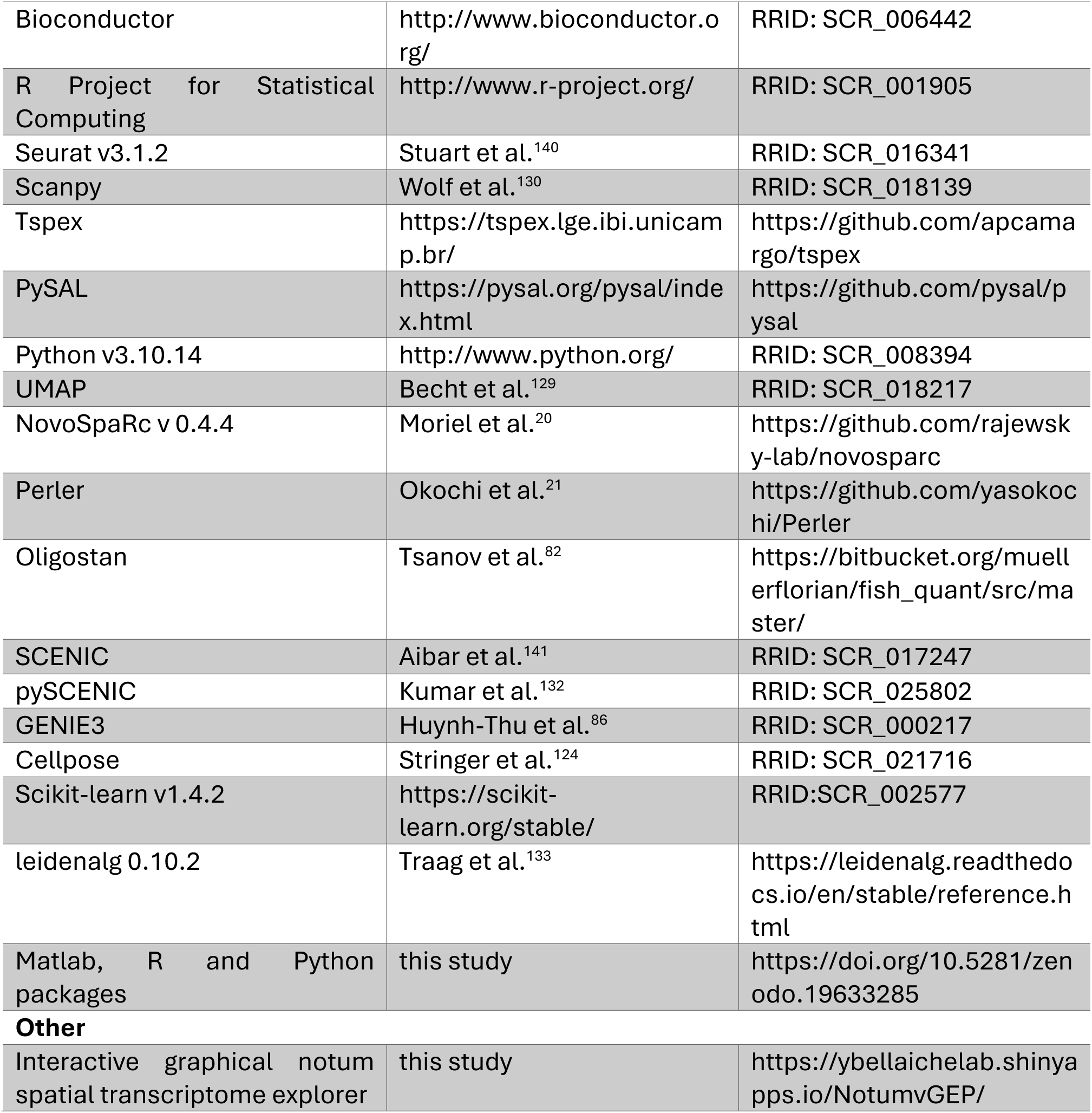

## Supplementary Figure Legends

**Figure S1.**
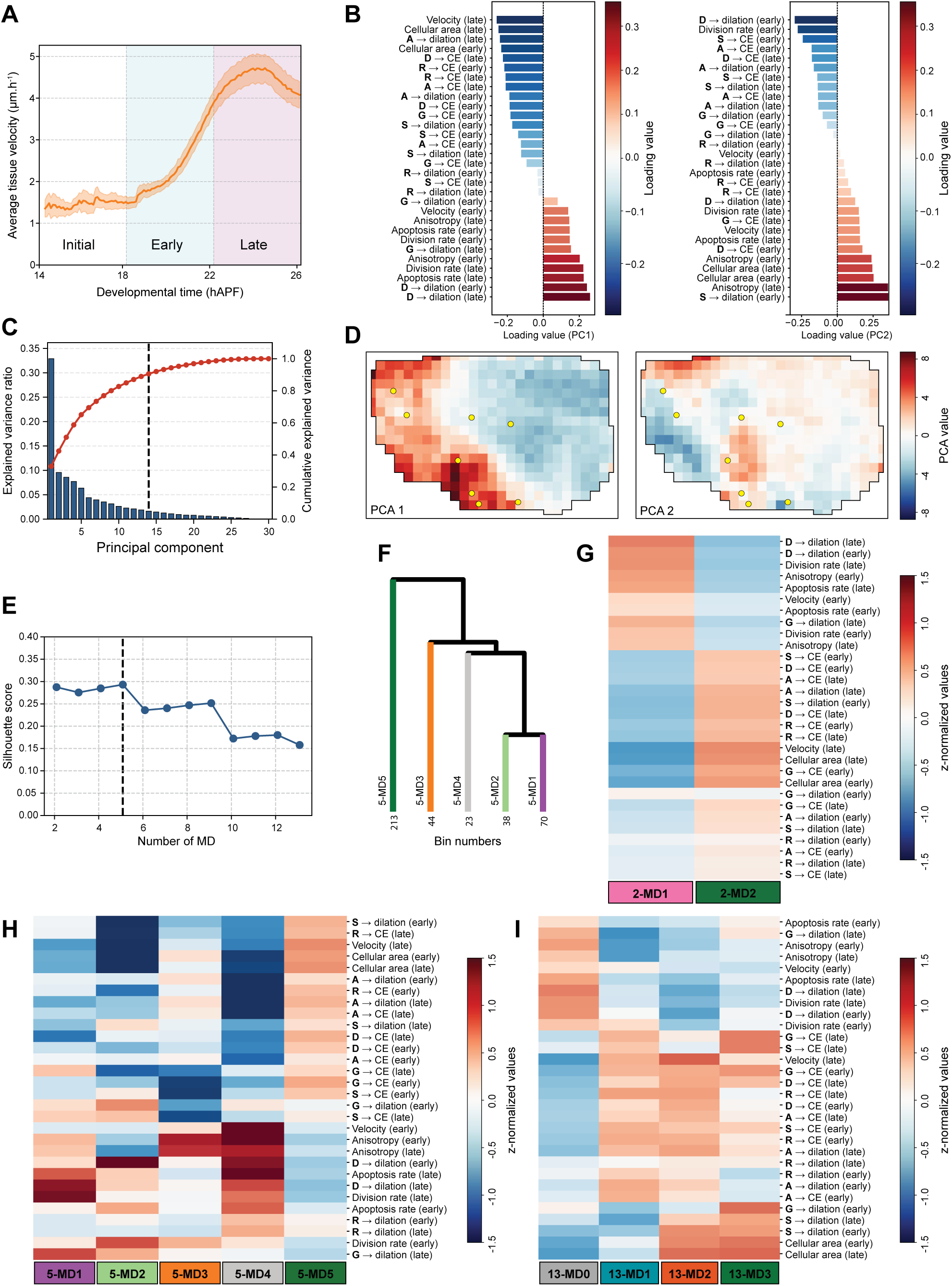
Hierarchical clustering of tissue dynamics in morphodynamic domains, related to Figure 1. (**A**) Global average tissue velocity magnitude (± SEM) over developmental time for *N* = 8 live image movies. Background colors mark three phases: slow initial phase (14-18 hAPF), acceleration early phase (18-22 hAPF), and late fast phase (22-26 hAPF). (**B**) Loading values of the 15 features in the early and late phase, indicating their contributions to the principal components PC1 (left) and PC2 (right). ‘**X** → dilation’: contribution of a cellular process to tissue isotropic dilation. ‘**X** → CE’: contribution of a cellular process to tissue convergent extension (EC) (see Table S1). (**C**) Bar plot of the percentage of variance explained by each principal component with the cumulative explained variance (red line). Vertical black dashed line indicates the number of components used for spatial clustering (14). (**D**) Spatial projection of principal component values for PC1 and PC2. Yellow dots: landmark macrochaetae. (**E**) Graph of the silhouette score as a function of the number of morphodynamic domains. Dashed line: optimal score of 0.259 obtained for 5-MD. (**F**) Dendrogram of hierarchical clustering showing cuts for the 5-MD morphodynamic domains. The number of spatial bins for each cluster, reflecting cluster size, is indicated. (**G-I**) Heatmaps of the z-normalized values of 15 features in the early and late phase for 2-MD (G), 5-MD (H), and the three anterior sub-clusters for 13-MD, alongside the background aggregated domain 13-MD0 (I). ‘**X** → dilation’: contribution of a cellular process to tissue isotropic dilation. ‘**X** → CE’: contribution of a cellular process to tissue convergent extension (CE) (see Table S1).

**Figure S2.**
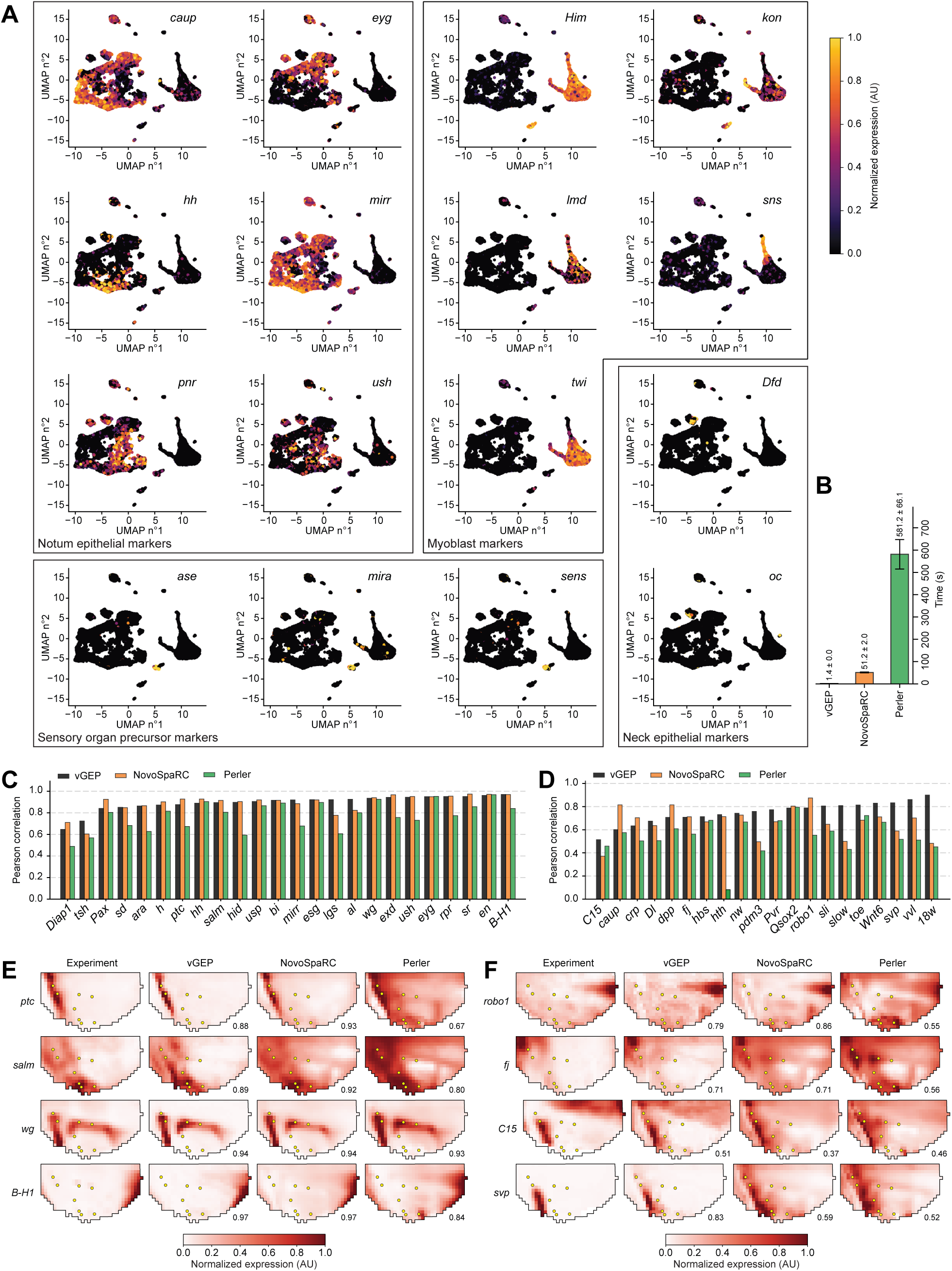
scRNA-seq data quality control and annotation, related to Figure 2. (**A**) UMAP projections of *n* = 12,187 single cells. Colors indicate expression levels of marker genes used to annotate cell type populations: Notum epithelial markers (*caup*, *eyg*, *hh*, *mirr*, *pnr*, *ush*), Myoblast markers (*Him*, *kon*, *Imd*, *sns*, *twi*), Sensory organ precursor markers (*ase*, *mira*, *sens*), and Neck epithelial markers (*Dfd*, *oc*). (**B**) Bar plot showing the mean raw reconstruction time (± SD) for vGEP, NoveSpaRC, and Perler for *n* = 10 runs. (**C**) Bar plot of individual Pearson correlations between predicted and reference (atlas) spatial expression profiles for the 24 atlas genes (training set) using vGEP, NovoSpaRC, and Perler. (**D**) Bar plot of individual Pearson correlations for 20 validation gene expression patterns using vGEP, NovoSpaRC, and Perler. (**E**) Average experimental expression patterns and reconstructed profiles for 4 representative atlas genes (*ptc*, *salm*, *wg*, *B-H1*), obtained using vGEP, NovoSpaRC, and Perler. r values (Pearson correlation between experiment and reconstruction method) are indicated. Yellow dots: landmark macrochaetae. (**F**) Average experimental expression patterns and reconstructed profiles for 4 representative validation genes, smiFISH (*robo1*, *fj*), and protein (*C15*, *svp*), obtained using vGEP, NovoSpaRC, and Perler. r values (Pearson correlation between experiment and reconstruction method) are indicated. Yellow dots: landmark macrochaetae.

**Figure S3.**
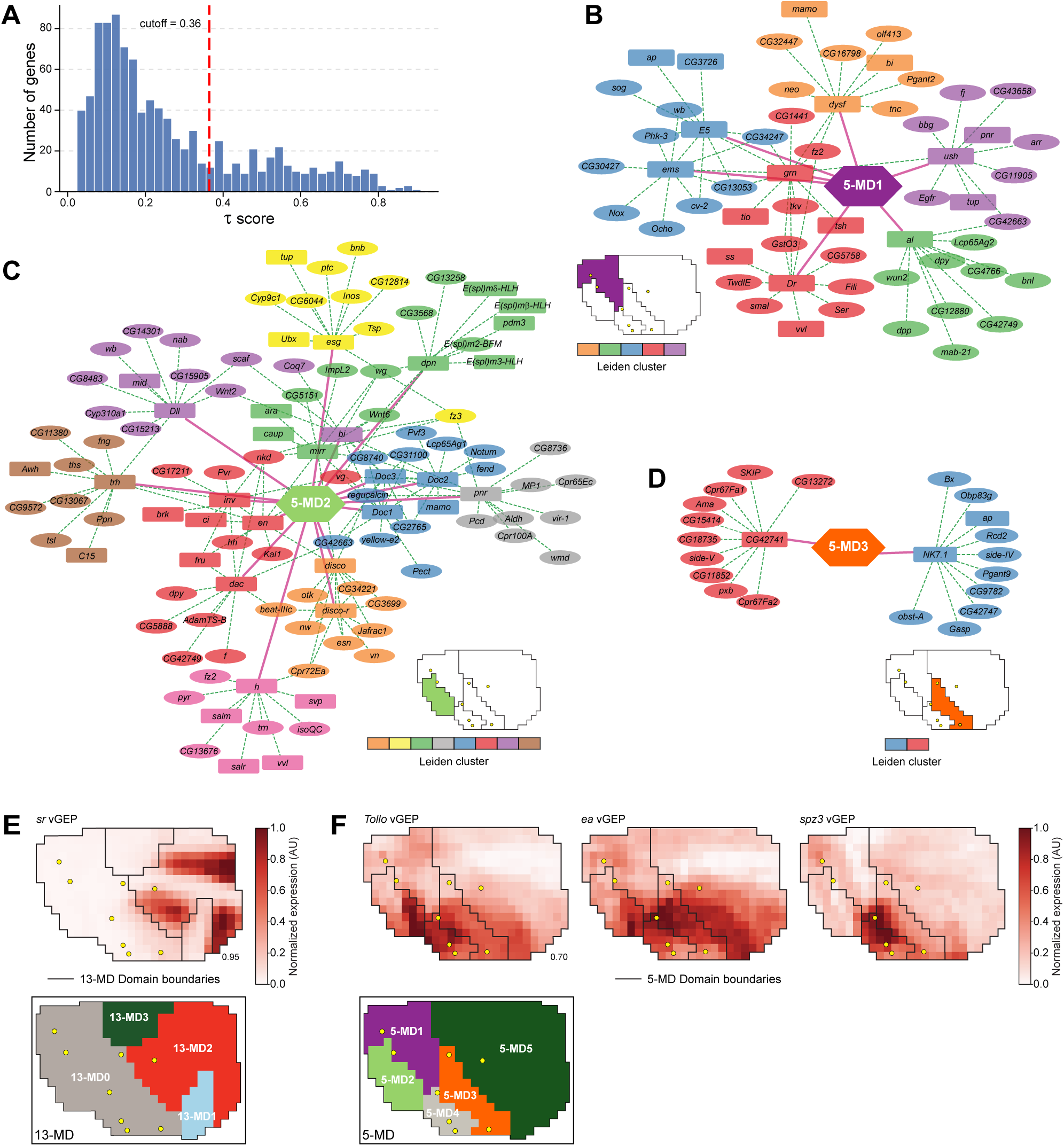
GRN analyses across 5-MD morphodynamic domains, related to Figure 3. (**A**) Distribution of tau scores for all genes maximally expressed in any of the 5-MD morphodynamic domains (*n* = 992 genes). Red dashed line indicates the specificity cutoff (τ > 0.36, *n* = 259 genes) used for downstream analysis based on Otsu’s thresholding. (**B-D**) Inferred GRNs for 5-MD1 (B), 5-MD2 (C), and 5-MD3 (D). Nodes represent TFs (rectangles) and their top 10 predicted target genes (ellipses). Colors indicate Leiden clusters. Pink solid lines indicate high specificity (τ > 0.36). Green dashed lines indicate putative regulation identified through GRN inference. (**E**) The predicted vGEP expression profile of *sr* is highly specific to 13-MD2. Black lines delineate the 13-MD boundaries. r value (Pearson correlation between experiment and vGEP) is indicated. The spatial map of 13-MD is indicated at the bottom. Only the anterior domain resolved into three sub-clusters is shown, whereas the posterior domain is shown in grey as a single aggregated cluster (13-MD0). Yellow dots: landmark macrochaetae. (**F**) Predicted vGEP expression profiles of *Tollo*, *ea* and *spz3*. The r value (Pearson correlation between experiment and vGEP) for *Tollo* is indicated. Black lines delineate the 5-MD boundaries. The spatial map of 5-MD is indicated at the bottom. Yellow dots: landmark macrochaetae.

**Figure S4.**
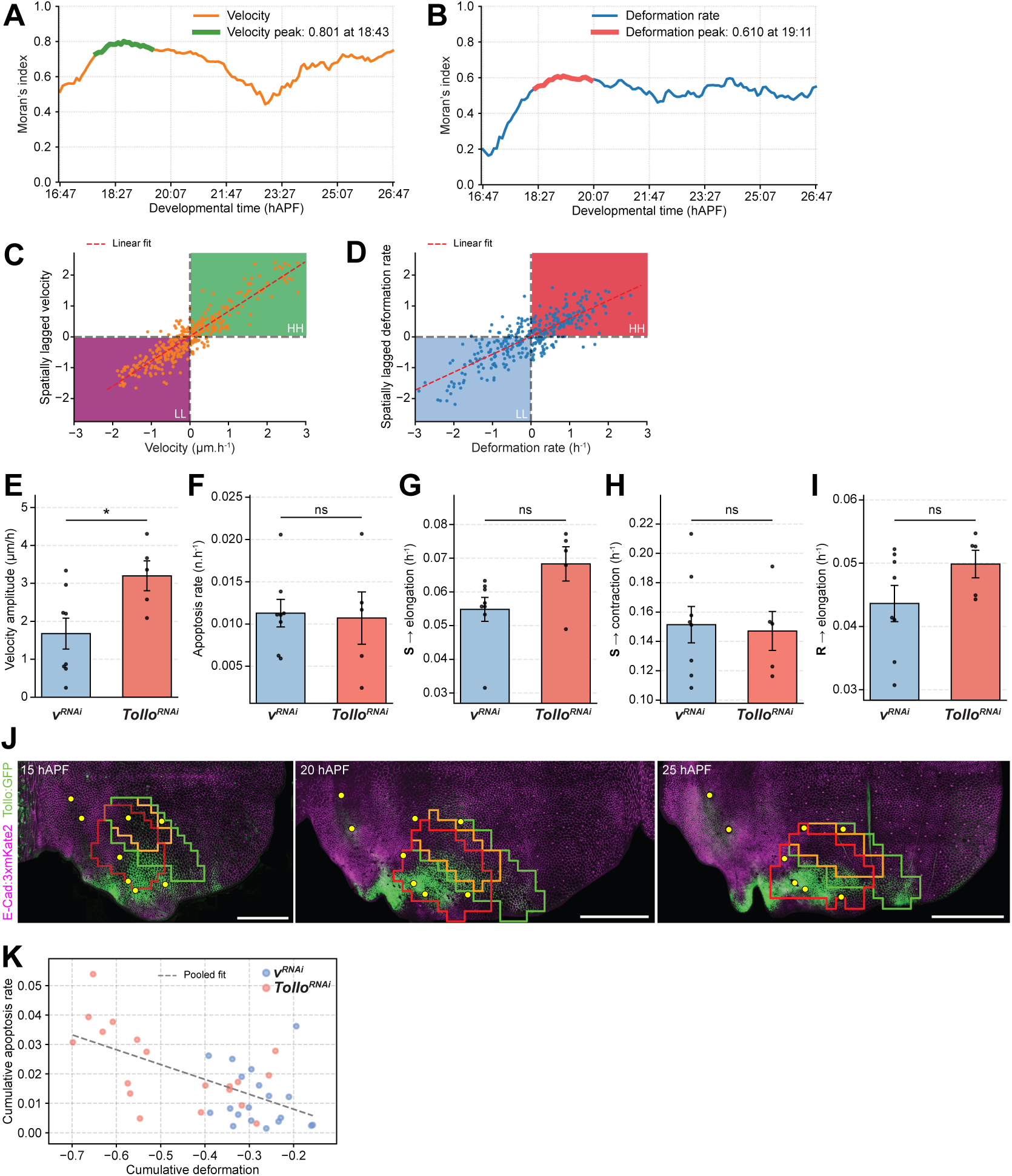
Moran’s index statistics on velocity and deformation profile differences, related to **Figure 4**. (**A**) Global Moran’s index for velocity differences between *Tollo^RNAi^* (*N* = 5) and control *v^RNAi^* (*N* = 8 animals) (*Tollo^RNAi^*-*v^RNAi^*). (**B**) Global Moran’s index for deformation rate differences between *Tollo^RNAi^* (*N* = 5) and control *v^RNAi^* (*N* = 8) (*Tollo^RNAi^*-*v^RNAi^*). (**C**) Correlation between velocity and spatially lagged velocity, defined as the mean value over its 8 queen-contiguity neighbors in a 2 h sliding window around the peak of Moran’s index (18:43 hAPF, see A) for each spatial bin. Pearson correlation coefficient *r* = 0.92. Red dashed line: linear regression fit (ordinary least squares). Two-sided t-test on the regression slope (*p* < 1 × 10^−@^). Values are sorted into 2 quadrants: LL (purple) corresponding to lower-than-average velocity values associated to lower-than-average velocity in the neighboring regions. HH (green) corresponding to higher-than-average velocity values associated to higher-than-average velocity in the neighboring regions. (**D**) Correlation between deformation rate and spatially lagged deformation, defined as the mean value over its 8 queen-contiguity neighbors rate in a 2 h sliding window around the peak of Moran’s index (19:11 hAPF, see B) for each spatial bin. Pearson correlation coefficient *r* = 0.85. Red dashed line: linear regression fit (ordinary least squares). Two-sided t-test on the regression slope (*p* < 1 × 10^−@^). Values are sorted into 2 quadrants: LL (blue) corresponding to lower-than-average dilation values associated to lower-than-average dilation in the neighboring regions. HH (red) corresponding to higher-than-average dilation values associated to higher-than-average dilation in the neighboring regions. (**E**) Comparison of the mean flow velocity amplitude (± SEM) between control *v^RNAi^* (*N* = 8) and *Tollo^RNAi^* (*N* = 5) tissues in the red ROI defined in Figure 4C, using a 2 h sliding window around the peak of Moran’s index in Figure S4A. Two-sided Mann-Whitney U test (\**p* < 0.05). (**F**) Comparison of the mean apoptosis rate (± SEM) between control *v^RNAi^* (*N* = 8) and *Tollo^RNAi^* (*N* = 5) tissues in the red ROI defined in Figure 4C, using a 2 h sliding window around the peak of Moran’s index in Figure S4A. Two-sided Mann-Whitney U test (ns, not significant). (G) Comparison of the mean contribution of cell shape changes to CE (± SEM) between control *v^RNAi^* (*N* = 8) and *Tollo^RNAi^* (*N* = 5) tissues in the red ROI defined in Figure 4C, using a 2 h sliding window around the peak of Moran’s index in Figure S4A. Two-sided Mann-Whitney U test (ns, not significant). (**H**) Comparison of the mean contribution of cell shape changes to tissue contraction (± SEM) between control *v^RNAi^* (*N* = 8) and *Tollo^RNAi^* (*N* = 5) tissues in the red ROI defined in Figure 4C, using a 2 h sliding window around the peak of Moran’s index in Figure S4A. Two-sided Mann-Whitney U test (ns, not significant). (**I**) Comparison of the mean contribution of rearrangements to CE between control *v^RNAi^* (*N* = 8) and *Tollo^RNAi^* (*N* = 5) tissues in the red ROI defined in Figure 4C, using a 2 h sliding window around the peak of Moran’s index in Figure S4A. Two-sided Mann-Whitney U test (ns, not significant). (**J**) E-Cad:3xmKate2 and Tollo:GFP confocal images showing Tollo:GFP expression in the notum at 15, 20 and 25 hAPF. The flow ROI (red, Figure 4C), contraction ROI (green, Figure 4D) and restricted contraction ROI (orange, Figure 4D) are rescaled to the reference archetype and plotted onto the individual images. Yellow dots: landmark macrochaetae. (**K**) Correlation between total contraction over 16:45–26:45 hAPF and total delamination in control *v^RNAi^* (*N* = 8) and *Tollo^RNAi^* (*N* = 5) tissues within each spatial bin in the orange ROI defined in Figure 4D. Pearson correlation coefficient *r* = 0.6. Grey dashed line: linear regression fit across both populations. Two-sided t-test on the regression slope (*p* < 1 × 10^−@^). Scale bars: 100 µm (J).

## Supplemental video titles and legends

**Video S1, related to Figure 1. Tissue velocity flow and deformation rate of on E-Cad:3xGFP hemi-notum from 13:57 to 25:54 hAPF.**

A. E-Cad:3xGFP hemi-notum time-lapse movie from 13:57 to 25:54 hAPF.
B. Movie of the tissue flow velocity measured on the time-lapse movie shown in A. Orange arrows indicate the direction and amplitude of tissue flow.
C. Movie of the tissue deformation rate computed from the flow velocity measured in B. Grey circles indicate the amplitude of isotropic contraction, and white circles indicate the amplitude of isotropic dilation, while bars indicate the orientation and anisotropy of CE rate.

**Video S2, related to Figure 1. Tissue morphometric measurements during pupal notum development from 13:57 to 25:5G hAPF.**

(**A**-**H**) Movies of the average (*N* = 8) tissue flow velocity (A), tissue deformation (B), cell area (C), cell anisotropy (D), as well as the contributions of cell shape change **S** (E), of apoptosis **A** (F), of cell division **D** (G), and rearrangements **R** (H) to tissue deformation. Cell area and cell anisotropy are represented as circle; tissue flow velocity by arrows corresponding to the direction and amplitude of tissue flow; deformation rates by grey circles indicating the amplitude of isotropic contraction, white circles indicating the amplitude of isotropic dilation, and bars indicating the orientation and anisotropy of CE rate.

**Video S3, related to Figure 4. Characterization of differences in velocity flow and isotropic deformation rate between *Tollo^RNAi^* and *v^RNAi^* tissues.**

(**A**, **C**, **E**) Movies of the average tissue flow velocity fields of the difference between *Tollo^RNAi^* (*N* = 5) and control *v^RNAi^* (*N* = 8) (*Tollo^RNAi^* - *v^RNAi^*) between 17:59–25:46 hAPF. Orange arrows indicate the direction and amplitude of tissue flow. In A tissue flow velocity fields are plotted over the local indicators of spatial association (LISA) based on local Moran’s index statistic. Shadings indicate LISA based on local Moran’s index statistic. Green shading denotes “High-High” (HH) spatial bins representing regions of significantly increased velocity difference; purple shading denotes “Low-Low” (LL) spatial bins of significantly decreased velocity difference. Color intensity reflects statistical significance as − log10(*p*). In C tissue flow velocity fields are plotted over the advected average Tollo:GFP expression pattern (*N* = 9). In E, tissue flow velocity fields are plotted over the map of the 5-MD morphodynamic domains advected onto a reference animal.

(**B**, **D**, **F**) Movies of the average tissue isotropic deformation rate fields of the difference between *Tollo^RNAi^* (*N* = 5) and control *v^RNAi^* (*N* = 8) (*Tollo^RNAi^* - *v^RNAi^*) between 17:59–25:46 hAPF. Grey circles indicate the amplitude of isotropic contraction; white circles indicate the amplitude of isotropic dilation. In B tissue dilation rate fields are plotted over the local indicators of spatial association (LISA) based on local Moran’s index statistic. Shadings indicate LISA based on local Moran’s index statistic. Red shading denotes “High-High” (HH) spatial bins representing regions of significantly increased tissue dilation; blue shading denotes “Low-Low” (LL) spatial bins representing regions of significantly increased tissue contraction. Color intensity reflects statistical significance as − log10(*p*). In D, tissue isotropic deformation rate fields are plotted over the advected average Tollo:GFP expression pattern (*N* = 9). In F, tissue isotropic deformation rate fields are plotted over the map of the 5-MD morphodynamic domains advected onto a reference animal.

