## Supplementary material for "Morphodynamic domains enable integration of live morphometrics and spatial transcriptomics": Sup Table 1

**Table S1. Quantitative descriptors used to characterized epithelial morphogenesis.**

| Feature | Description | Unit |
| --- | --- | --- |
| G → dilation | Tissue isotropic dilation rate | h⁻¹ |
| D → dilation | Division contribution to tissue isotropic dilation rate | h⁻¹ |
| R → dilation | Cell rearrangement contribution to tissue isotropic dilation rate | h⁻¹ |
| S → dilation | Cell shape change contribution to tissue isotropic dilation rate | h⁻¹ |
| A → dilation | Apoptosis contribution to tissue isotropic dilation rate | h⁻¹ |
| G → CE | Tissue convergent extension rate (deviatoric part of the tissue deformation tensor) | h⁻¹ |
| D → CE | Division contribution to tissue convergent extension rate | h⁻¹ |
| R → CE | Cell rearrangement contribution to tissue convergent extension rate | h⁻¹ |
| S → CE | Cell shape change contribution to tissue convergent extension rate | h⁻¹ |
| A → CE | Apoptosis contribution to tissue convergent extension rate | h⁻¹ |
| Velocity | Tissue velocity | µm·h⁻¹ |
| Normalized division rate | Division rate normalized by the initial number of cells | Dimensionless |
| Normalized apoptosis rate | Apoptosis rate normalized by the initial number of cells | Dimensionless |
| Anisotropy | Cell anisotropy | Dimensionless |
| Cellular area | Cell area | µm² |
